# Vegetation of *Chamaecyparis* montane cloud forest in Lalashan Forest Dynamics Plot

**DOI:** 10.1101/2023.11.04.565604

**Authors:** Ting Chen, Yi-Nuo Lee, Po-Yu Lin, Kun-Sung Wu, David Zelený

## Abstract

To understand the vegetation-environment relationships within the *Chamaecyparis* montane mixed cloud forest in Taiwan, we established a 1-ha Lalashan Forest Dynamics Plot (LFDP) in northern Taiwan (24°42D N, 121°26D E). We established the plot in July 2019 and finished the first census of all woody species in August 2020. We collected environmental factors related to the topography and soil properties, measured microclimate and soil moisture within the plot, and collected further microclimatic data with a nearby weather station. In total, we recorded 5220 individuals belonging to 65 species, 42 genera and 29 families, with a basal area of 69.1 m^2^ ha^−1^, dominated by *Chamaecyparis obtusa* var. *formosana*, *Rhododendron formosanum* and *Quercus sessilifolia*. Modified TWINSPAN classified vegetation into three types (ridge, east-facing slope and valley). Unconstrained ordination showed that the main gradients behind compositional changes are related to windwardness and convexity. The prevailing wind direction in the area is from the northeast, linked to the winter monsoon. Both east-facing slope type and valley type have relatively lower temperatures than ridge type, especially during summer. Convexity is related to soil moisture gradient (from dryer convex to wet concave sites). Fog frequency is seasonal, with the highest values during autumn and winter months. From soil properties, pH is negatively and phosphorus is positively related to topographical convexity. Litter decomposition is linked to both topographical, soil and biotic variables. Collected data will serve as a baseline for future resurveys and monitoring changes within this montane cloud forest.

## Introduction

Montane cloud forests (MCFs) are characterised by the presence of persistent and frequent wind-driven cloud and foggy conditions at the ground (tree) level (Hamilton 1995). The distribution of montane cloud forests is highly fragmented according to the restriction of persistently foggy zones, making these fragments act like isolated islands which are assumed to promote speciation and endemism (Bruijnzeel et al. 2010; Li et al. 2015). MCFs are one of the world’s most endangered ecosystems because of their sensitivity to changes in unique ecological conditions (Bruijnzeel et al. 2010). In addition, climate observations show that MCFs are suffering from a decreasing trend in ground fog occurrence which is likely related to climate change (Still et al. 1999; Foster 2001; Ponco-Reyes et al. 2012; Hu and Riveros-Iregui 2016).

High fog frequency in montane cloud forests is responsible for the occurrence of special environmental conditions that are different from other forest types, including horizontal precipitation, high air humidity, lower light availability, lower air temperature, and chronic nutrient limitation in soil (Stadtmüller 1987). Horizontal precipitation represents an extra water input, in addition to rainfall (vertical precipitation), and is formed when fog condensates on leaf surfaces, a process also known as "fog stripping" (Stadtmüller 1987). High air humidity mitigates the temperature differences, however, it hinders the leaf transpiration process and makes epiphylls (including lichens, mosses, algae, and fungi) grow on and cover the leaves more easily, which may result in reducing photosynthesis (Lai et al. 2006). The presence of fog can reduce 10–15% of light compared to no-fog conditions, causing lower light availability for plants, and hence, lower photosynthetic rate, but may lower the effect of photoinhibition and increase photosynthesis efficiency under diffuse light at the same time (Urban et al. 2007; Reinhardt and Smith 2008). When fog occurs, the air temperature is 3–6°C lower than analogous site without fog, alleviating possible heat stress for plants, but the plants may encounter frost events. Lower air temperature may also lead to relatively low overall heat income for plants, and thus decreases the efficiency of photosynthesis (Lai et al. 2006). Due to very frequent high air/soil humidity and lower air temperature, the decomposition rates in montane cloud forests are slower, causing chronic nutrient limitation in soil (Tanner et al. 1990).

Past studies about montane cloud forests were mostly focusing on tropical regions, while fewer studies were done in subtropical montane cloud forests (SMCF), although its biological and conservation value is not less significant (Li et al. 2015). In subtropical eastern Asia, a large proportion of montane cloud forests are evergreen broadleaved forests mixed with coniferous and deciduous broadleaved trees (Su 1984; Li et al. 2015). By a common dominance of coniferous species, the subtropical montane cloud forests differ from tropical ones, which are dominated only by evergreen broad-leaved trees (Bruijnzeel et al. 2010).

Zonal forests in Taiwan can be classified into five vegetative zones based on local climate, which is primarily driven by elevation, and at elevations around 1500 to 2500 m a.s.l., the montane zone is characterised by frequent ground fog occurrence (Li et al. 2015; Schulz et al. 2017). In some areas, montane cloud forests distribute in lower elevations than 1500 m a.s.l., partly due to the influence of the northeastern monsoon, which locally decreases the temperature, especially in the northeastern part of Taiwan (Lai et al. 2006; Li et al. 2013, 2015; Schulz et al. 2017), and also as the result of the mass elevation heating effect (Massenerhebung effect, Quervain et al. 1904), which decreases the elevation of vegetation zones on smaller and isolated mountains as a result of lower heating effect (Su 1984).

Montane cloud forests in Taiwan include three main subtropical vegetation types, namely *Chamaecyparis* montane mixed cloud forest, *Fagus* montane deciduous broad-leaved cloud forest, and *Quercus* montane evergreen broad-leaved cloud forest, and one tropical vegetation type, namely *Pasania-Elaeocarpus* montane evergreen broad-leaved cloud forest (Li et al. 2013).

Detailed forest dynamics plot (FDP) studies allow us to better understand the forest ecosystems, and with long-term plot resurveys, we can also monitor vegetation dynamics in the sense of species composition changes. This is essential not only for improving theoretical knowledge about the dynamics of the montane cloud forest vegetation but eventually also to describe their response to ongoing climate change. However, there are only three previous FDP studies in Taiwan that are focused on SMCF, one in Yuanyang Lake Long-Term Ecological Research Site (Chou et al. 2000) and two in Mt. Peitungyen in central Taiwan (Song 1996; Song et al. 2010; Hu and Tzeng 2019); none of them however studied also the effect of northeastern monsoon. On the contrary, previous FDP studies focused on the effects of monsoon on forest vegetation in Taiwan are mostly below the cloud forest zone, namely one FDP in the submontane rainforest at Lopeishan in northern Taiwan (Lin et al. 2005) and four FDPs in lowland subtropical rainforest at Nanjenshan region in southern Taiwan (Chao et al. 2007, 2010). Studies done in Nanjenshan and Lanjenchi forest dynamics plots found that forests under different wind exposure differ in species composition, and that windward forests have denser, shorter, and smaller trees than the leeward forests (Ku et al. 2021).

Our study was done in the Lalashan Forest Dynamics Plot (LFDP), established in a SMCF in the northern part of Taiwan, on a flat ridge with east-facing slope influenced by the northeastern monsoon. The vegetation of LFDP belongs to *Chamaecyparis* montane mixed cloud forest, where coniferous and broad-leaved woody species co-occur, with an admixture of deciduous species (Li et al. 2015). We established LFDP in 2019, in order to learn more about the vegetation and the environmental factors influencing a SMCF, and to collect baseline data to monitor the future vegetation changes. The aims of our study are: 1) to distinguish main vegetation types and describe their compositional and environmental differences; 2) to describe the main gradients of changes in woody species composition within the permanent plot, and the possible mechanism; and 3) to explore environmental conditions of the plot, including soil chemical properties, litter decomposition and microclimatic conditions.

## Materials and Methods

### Study site

Lalashan Forest Dynamics Plot (LFDP; 24°42D N, 121°26D E; elevation 1758–1782 m a.s.l.) is located on a wide part of the mountain ridge near the saddle between Lalashan and Tamanshan, inside the Chatianshan Nature Reserve, in northern Taiwan (Fig. 1). The mountain ridge is in the northern branch of Xueshan Range, with orientation of northwest-southeast direction. There are two west-east direction ephemeral streams in the western part of LFDP, and an east-facing slope exposed to the northeastern monsoon in the eastern part of LFDP. The vegetation in LFDP is *Chamaecyparis* montane mixed cloud forest (Li et al. 2013), which is dominated by *Chamaecyparis obtusa* var. *formosana* (coniferous species) and *Rhododendron formosanum* (evergreen broadleaf species).

**Figure 1.**
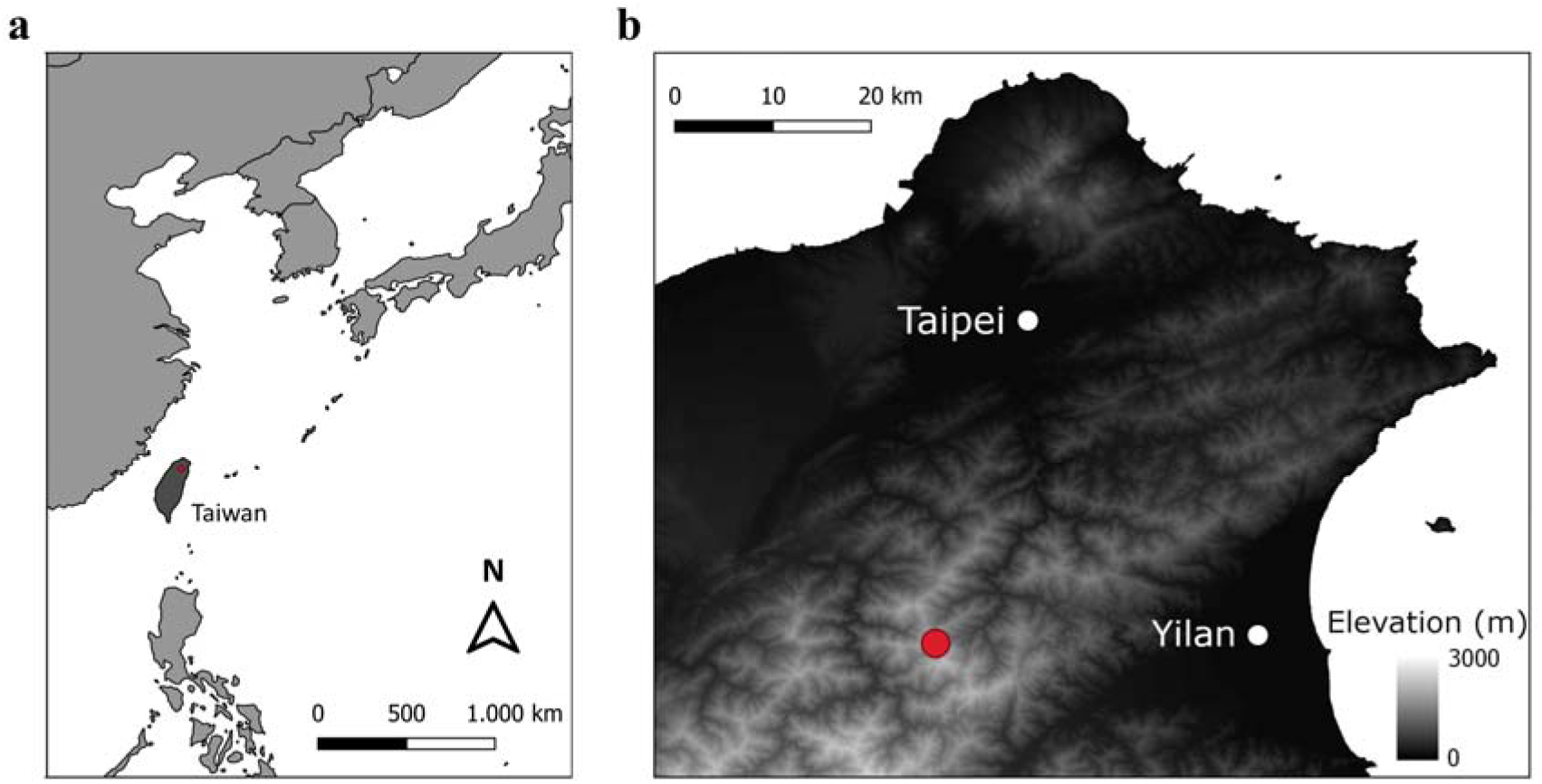
Location of LFDP (red point) in East Asia (a) and in northern Taiwan (b)

### Sampling design

We established the boundaries of the plot and all subplots in July 2019, and finished the first census of woody species in August 2020. The establishment and survey of woody species followed the Forest Global Earth Observatory Network (ForestGEO) Tree Census Protocol (Condit 1998). We used the compass with a telescope (Ushikata LS-25, Kantum Ushikata Co. Ltd., Yokohama, Japan) to delineate a projected area of one hectare (100 m × 100 m), which was then subdivided into 100 10 m × 10 m subplots (Fig. 2). The aspect of the main LFDP axis is pointing to the north, and the corners of the subplots were coded with the coordinates, with (0,0) starting from the southwest corner and ending with (10,10) in the northeast corner. The corners of the subplots were marked with PVC poles painted red at the top, and the centres of the subplots were marked with plastic poles painted yellow at the top. The subplot IDs were coded according to the coordinates of their southwest corners.

**Figure 2.**
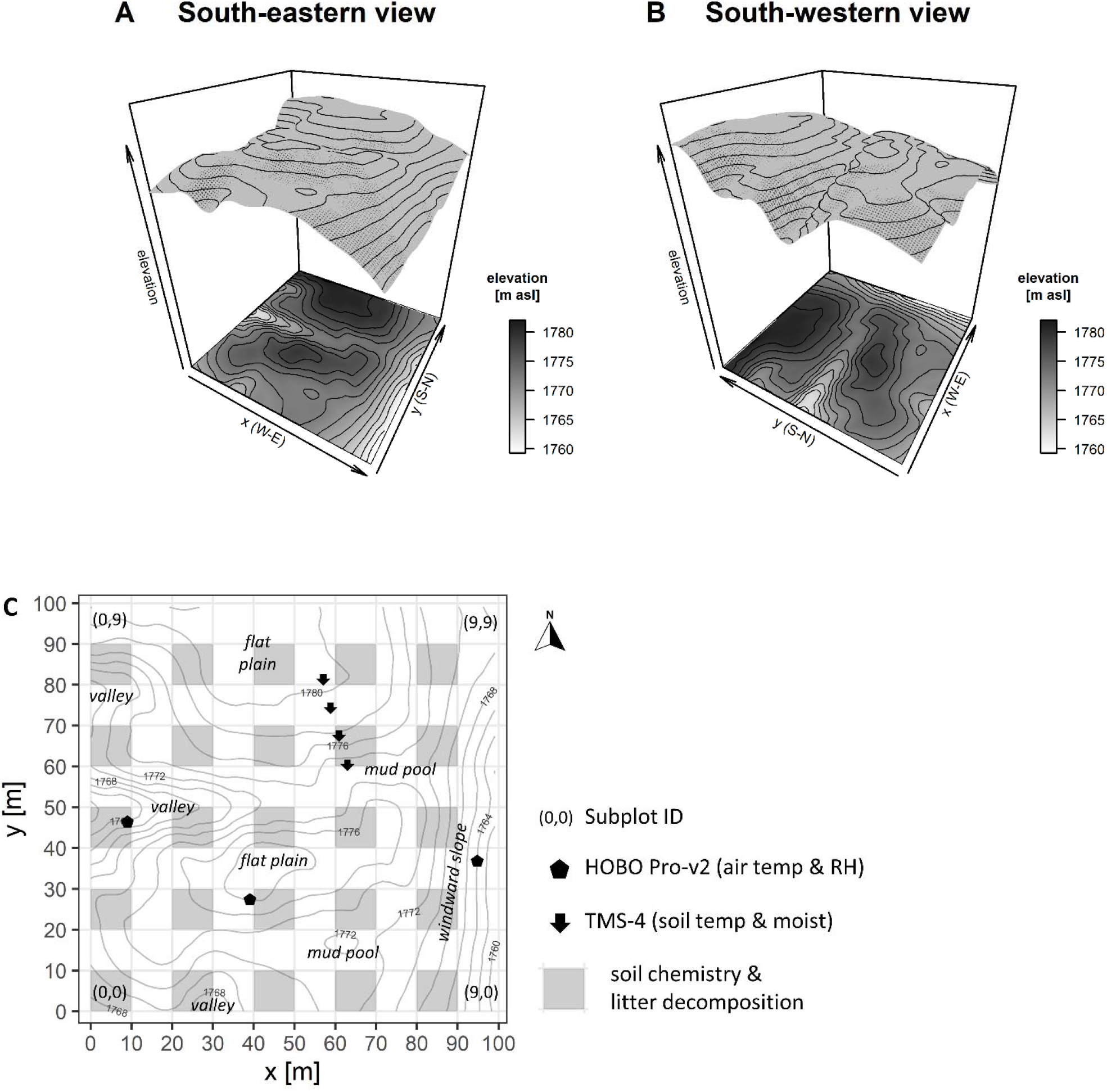
Maps of the Lalashan Forest Dynamics Plot. Panels A and B show a 3D topographical model from two views (south-eastern and south-western, respectively). Panel C shows a 2D map with the positions of HOBO Pro-v2 loggers (air temperature and relative humidity, RH), TMS-4 loggers (soil temperature and moisture), and subplots with measured soil chemistry and litter decomposition. Text labels highlight typical topographical features of the plot.

### Species composition

When surveying, we first delineated the boundaries of each subplot with a tape, and only the trees rooted inside the surveyed subplot were recorded for the subplot. In each subplot, all individuals of woody species (excluding lianas) with diameter at breast height (DBH) ≥ 1 cm were identified, tagged with iron tag with a stamped number, mapped, and their DBH measured. If the individual tree had any branches with DBH ≥ 1 cm, we also measured them and tagged them with white plastic tag with a number written by a pencil.

For each species, we calculated importance value index (IVI; Curtis 1959) as the sum of relative basal area (BA) and relative individual density of each species in the subplot. Basal area of both main stem and all branches were considered; for individual density, only the main stem was considered as individual.

For each subplot, we calculated the density of individuals (as the number of all main stems in the subplot), total BA (as the sum of BA of all species in the subplot), maximal BA (the BA of the tree with highest DBH), and the mean number of branches per individual. The species in LFDP were categorised into three leaf types including conifer, deciduous broadleaf and evergreen broadleaf species, using information from Flora of Taiwan, 2^nd^ edition (Huang & Hsieh 1994–2003) and our field observations. Using leaf type category, for each subplot we calculated BA of all conifer species, all deciduous species and all evergreen broadleaf species.

### Topographical variables

The environmental factors related to topography, including elevation, convexity, slope, aspect and variables calculated of them (such as northeasterness and windwardness) were all derived from the elevation of the poles in corners of subplots. We first measured the elevation of the pole (5,0) GPS (GARMIN GPSMAP 64st, USA), and elevations of all other poles were calculated using the slope angles between poles recorded while delineating the plot. Then, for each subplot, the elevation was calculated as the mean elevation of its four corner poles. The convexity was calculated as the elevation of given subplot minus the mean elevation of its eight-surrounding subplots; for marginal subplots it was calculated as the elevation of the subplot’s centre pole (additionally measured in the field with the compass and telescope) minus the elevation of the subplot (calculated as the mean elevation of the four corner poles). The slope was calculated as the mean angular deviation from the horizon of each of the four triangular planes formed by connecting three of the target subplot’s corner poles. The aspect was calculated as the elevation of the midpoints of each subplot’s four sides by averaging the elevation of the two corner poles on each side, using the formula 180 – arctan (fy / fx) · (180 / π) + 90 (fx / |fx|), where fx is the midpoint elevation change from the east side to west side, and fy is the midpoint elevation change from the north side to south side (Valencia *et al*. 2004, De Cáceres *et al*. 2012). For analysis, we converted aspect into northeasterness by folding it along the SW-NE direction (with 0° for SW and 180° for NE). The windwardness was calculated by multiplying the aspect, folded along the E-W axis (of prevailing wind) and centred around zero (by setting + 90° in the east and –90° in the west), by slope; the highest values are in steeper, east facing slopes that are the most wind-affected, while the lowest values are in steeper west facing slopes, which are shaded from the wind.

### Soil properties

Within all 100 subplots, we measured soil properties and collected soil samples for lab analysis. Within a subset of 25 subplots we made comprehensive physical and chemical analysis of soil samples, and also conducted teabag decomposition experiment. Each 10 m × 10 m subplot was divided into four 5 m × 5 m quadrats, and soil properties and soil samples were collected from centre of each quadrat. The soil properties measured in all 100 subplots include soil depth (measured with a 30 cm long iron rod, 0.6 cm in diameter), soil rockiness (estimated as the relative proportion of stones in the soil when taking the soil samples, with values from four estimates per subplot averaged into one value). Soil depth ranged from 0-30 cm, and soil depth deeper than 30 cm was recorded as 30+ cm; the measured values were converted into an ordinal scale (0 = 0 cm; 1 = 1–5 cm; 2 = 6–10 cm; 3 = 11–20 cm; 4 = 21–30 cm; 5 = 30+ cm), and median of the four values within each subplot was calculated to represent subplot-level soil depth. Four soil samples were collected within each subplot with iron shovel from the depth 0-10 cm after removing the surface litter, and mixed into one mixed soil sample. The soil chemical properties were measured in the laboratory only in 25 selected subplots (Fig. 2C), except soil pH that was measured for all 100 subplots. The collected soil samples were air-dried in the lab for several weeks and sieved through 2.0 mm sieve (2.0 mm laboratory test sieve, Endecotts Ltd, England). From all 100 samples we measured soil pH, using a glass electrode pH meter (LAQUA F-71, Horiba Ltd., Kyoto, Japan) in the solution of soil sample and deionised water in 1:2 ratio (10 g of soil and 20 ml of deionised water). All other soil properties were measured in the soil from a subset of 25 subplots. Soil texture (sand, silt and clay) was measured by hygrometer method (Gee & Bauder 1986); organic C content was acquired by Walkley and Black dichromate method (Nelson & Sommers 1996); total N was determined by Kjeldahl method (Nelson & Sommers 1972); C/N ratio was calculated as organic C divided by total N; exchangeable N, which contained the ammonium-N and nitrate-N, was determined by KCl extraction and steam distillation (Mulvaney 1996); available P was determined by Bray No. 1 method (modified from Burt, 2004) with a spectrometer (UV-1900PC, Macylab Instruments Inc., Shanghai, China); exchangeable cations of K, Ca and Mg were extracted by 1 M ammonium acetate (pH 7) and determined by a flame atomic absorbance spectrophotometer (AAnalyst 200, PerkinElmer, Inc., Waltham, MA, USA; Burt 2004); and available cations of Fe, Cu, Zn were extracted by 0.1 N HCl and determined by AAnalyst 200 (Baker & Amacher 1982).

Decomposition rate and stabilisation factor were also measured for the 25 subplots, following the protocol proposed by Keuskamp et al. (2013). Four pairs of green tea and rooibos tea commercial teabags (Lipton, EAN: 8714100770542 and 8722700188438) were buried 8 cm deep in the soil in the 25 subplots, and after 102–104 days (between July 13–15 and October 25– 26, 2020), the teabags were collected back to the lab. The teabags were first dried in oven at 70°C for 48 hours. Then, the remnants of tea were extracted from the plastic bag, weighted, and further combusted in a muffle oven at 550°C for 16 hours. Finally, the weights of the remnants subtracted the weights of the remains after combustion and calculated decomposition rate *k* and stabilisation factor *S* by formulas from Keuskamp et al. (2013).

### Microclimatic measurements

To quantify air and soil microclimatic parameters of the studied forest, we 1) renewed the standard weather station in the saddle near the forest dynamics plot, 2) installed air temperature and relative humidity loggers within the dynamics plot to describe differences between the vegetation types, and 3) installed soil moisture loggers within the dynamics plot along the convex-concave topographical gradient. The standard weather station is located in a small deforested saddle between Lalashan and Tamanshan mountain at elevation 1730 m a.s.l. (24°43’20.9"N 121°26’31.1"E). In the spring 2020, we used the tower (6.5 m tall) available from the previous station operated by Taiwan Forestry Bureau, and equipped it by new sensors and loggers. For the purpose of this study, we report long-term measurements of air temperature, relative humidity, precipitation, wind speed and direction, and visibility (as a proxy of fog intensity), measured from May 2020. Air temperature and relative humidity were first measured by ATMOS 14 4-in-1 sensor, housed in a sunshield, and after malfunction in December, 2021 by HOBO Pro-v2 logger, housed in a sunshield (installed at the the weather station in September 2021). Rainfall was measured by RS-102N Rain Gauge, wind speed and direction by Young 05103 WS/WD meter, and visibility by MiniOFS Sensor. The climatic data (except temperature and relative humidity recorded by HOBO logger) were computed and saved in the CR300-Series Datalogger every 10 minutes; HOBO Pro-v2 logger recorded the data every 30 minutes.

Within the dynamics plot, we installed three HOBO Pro-v2 loggers (housed in the sunshield), by attaching them on northern side of selected tree at height ca 1.4 m. The location was selected to measure microclimate in the three alternative topographical positions (Fig. 2C): valley in subplot (0,4), flat ridge in subplot (3,2), and east-facing wind-affected slope in subplot (9,3). Recording was conducted continuously from April 2020 with temperature and relative humidity recorded every 30 minutes. However, the sensor of the logger in the flat ridge was broken from October 2020 to June 2021.

Soil moisture and temperature were monitored by installing four TOMST TMS-4 loggers along the topographical gradient from the convex ridge to concave valley close to the centre of the dynamics plot (Fig. 2C). The measurement was done from April 17 till May 28, 2022, every 15 minutes, and included soil moisture and three values of temperature: 15 cm above soil surface, 2 cm above soil surface and 6 cm below the surface.

### Statistical analyses

Since some of the soil variables used for the analysis had skewed distribution, we transformed them by log_10_ transformation to improve their distribution (namely organic C, total N, C/N ratio, exchangeable N, available P, K, Ca, Mg, Cu and Zn). Pairwise correlations among pairs of soil variables was calculated using Spearmen’s rank correlation coefficient.

With subplot-based IVI data, we classified the forest vegetation at LFDP into three vegetation types by modified two-way indicator species analysis (modified TWINSPAN; Hill 1979; Roleček et al. 2009), using R package "twinspanR" (Zelený 2021). In modified TWINSPAN, pseudospecies cut levels were set to 0, 2, 5, 10, and 20%, and Bray-Curtis distance was used to measure compositional dissimilarity. For species composition differences between the three vegetation types, we determined diagnostic, dominant and constant species for each vegetation type, using JUICE software (Tichý 2002). Diagnostic species were determined as species with fidelity coefficient Φ ≥ 35 (Chytrý et al. 2002) in the subplots of a vegetation type, and significant at P < 0.05 when tested by Fisher’s exact test. Dominant species were determined as species with IVI ≥ 20%, and constant species were determined as species with frequency ≥ 80%. Each vegetation type was named by combination of the diagnostic species with the highest fidelity and the most dominant species for this vegetation type. Physiognomic and environmental differences between the three vegetation types by using analysis of variance (ANOVA) and Tukey’s honestly significant difference test (Tukey’s HSD).

To explore the main compositional gradients in species composition of the plot, we calculated detrended correspondence analysis (DCA; Hill and Gauch 1980) on IVI data transformed as log_10_ (x +1). We calculated multiple regression of each environmental variable on the first two DCA axes, including variables measured in all 100 subplots and those measured in the subset of 25 subplots, and projected them onto the ordination diagram if they were significant (at P < 0.05 for variables measured in all 100 subplots, and at P < 0.1 for those measured in 25 subplots). We also visually verified the relationships between the three vegetation types and environmental factors by projecting them onto the DCA ordination diagram. Relationship of environmental variables to DCA axes was calculated by *envfit* function in "vegan" package, version 2.5-7, Oksanen et al. 2020), with P-values calculated using toroidal permutation test to acknowledge the spatial autocorrelation (Legendre and Legendre 2012).

The DBH size-class analysis of each species was visualised using the histogram of the number of individuals, where the breaks for the DBH size classes were calculated using the following formula: M = INT (6*log_10_ N), R = (DBH_max_ – DBH_min_)/M, where N is the number of individuals for given species, DBH_max_ and DBH_min_ are the maximal and minimal DBH of given species, respectively, and R is the class-width parameter. The histogram was plotted only for species with at least 15 individuals. The final shape of the histogram was subjectively evaluated in terms of the regeneration status of the species.

Teabag decomposition data were analysed for their response to other measured topographical, soil and physiognomic variables. We took separately the decomposition rate *k* and stabilisation factor *S* as a response variable in the multiple linear regression model. As explanatory we then used topographical variables (elevation, slope, convexity, northeasterness and windwardness), soil variables (all including soil depth, rockiness, soil physical properties and chemical properties) and physiognomic variables (calculated for each subplot as total basal area of only conifer trees, only deciduous trees and only evergreen broadleaved trees). With all variables standardised to z-scores, we used forward selection to choose most parsimonious model for each litter decomposition variable (only variables significant at P < 0.05 during the forward selection were included in the final model).

For the microclimatic data analysis, we applied window-averaging algorithm with two-hour time interval on all temperature, relative air humidity and soil moisture measurements. In the weather station, the air temperatures (measured by ATMOS sensor from January 1, 2021 to December 29, 2021, and by HOBO Pro-v2 logger from December 30, 2021 to December 31, 2021) and the precipitation (measured from January 1, 2021 to December 31, 2021) were aggregated to draw climatic diagram. The visibility (measured from January 1, 2021 to December 31, 2023), was used to calculate fogginess and foggy time. The fogginess was classified into three categories, heavy, medium and light, when the visibility was less than 100 m, 100–500 m, and 500–1000 m, respectively. The foggy time was calculated as a percentage proportion of visibility measurements falling into the relevant category of fogginess out of all measurements done within a given month. The wind speed and wind direction (measured from January 1, 2021 to December 31, 2023) were aggregated to draw a wind rose chart. The air temperatures measured by HOBO Pro-v2 loggers within the LFDP from August 1, 2021 to July 31, 2023 were aggregated into mean monthly temperature, mean monthly daily temperature deviation, and the proportion of wet days (i.e. days with mean RH > 99%) and dry days (RH < 75%). The soil moisture measured from April 20 to May 20, 2022, was recalculated into mean daily soil moisture and volumetric water content.

All analyses in this study were done in R program version 4.3.0 (R Core Team 2023), with R code, vegetation data and environmental variables stored in the GitHub repository (https://github.com/zdealveindy/VegLab/tree/main/data/lalashan-fdp-vegetation).

## Results

A total of 5220 individuals belonging to 65 species, 42 genera and 29 families were recorded in LFPD, with a total BA of 69.1 m^2^ ha^−1^. The surveyed forest is dominated by *Chamaecyparis obtusa* var. *formosana* (14% of the whole-plot-based IVI), *Rhododendron formosanum* (14%), *Quercus sessilifolia* (9%), *Trochodendron aralioides* (7%) and *Eurya crenatifolia* (5%), with the cumulative IVI of these five most dominant species reaching 49%. Distribution maps drawn separately for coniferous, deciduous and evergreen broadleaf species (Fig. 3) show proportional differences between each leaf-habit type (with 4499 evergreen broadleaf, 447 deciduous, and 270 coniferous individuals), and also different patterns of their distribution. Conifers are distributed mainly on the flat plateaus while avoiding the valleys and steep east-facing slopes, deciduous species frequently occur on steep east-facing slopes and also extend to ridges, and evergreen broadleaf species are scattered across the whole plot, with visibly higher density in the eastern part. When considering individual 10 m × 10 m subplots (Fig. S3), species richness varied between 7 and 31 species, with median of 15; numbers of individuals varied between 11 and 184, with median of 41; sum of BA per subplot (recalculated per hectare) varied between 5.5 m^2^ ha^−1^ and 191.6 m^2^ ha^−1^, with average 69.1 m^2^ ha^−1^; maximal BA of the largest tree within the subplot varied between 138.9 cm^2^ and 8300.0 cm^2^, with average 2170.3 cm^2^; the mean number of branches per individual varied between 0.1 and 2.1, with average of 0.6.

**Figure 3.**
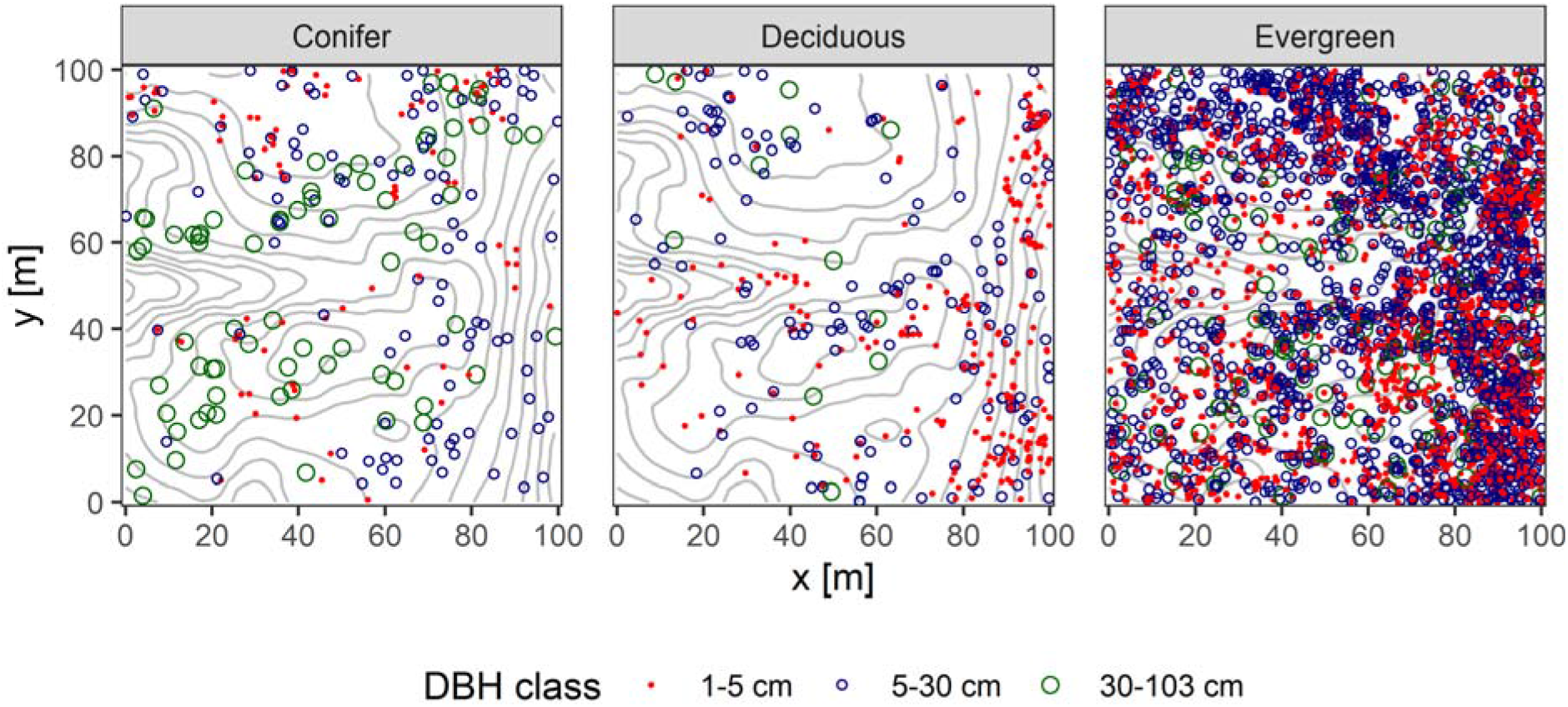
Distribution map of conifer, deciduous and evergreen tree individuals stratified into three DBH classes (1-5 cm, 5-30 cm, and 30-103 cm)

**Figure 4.**
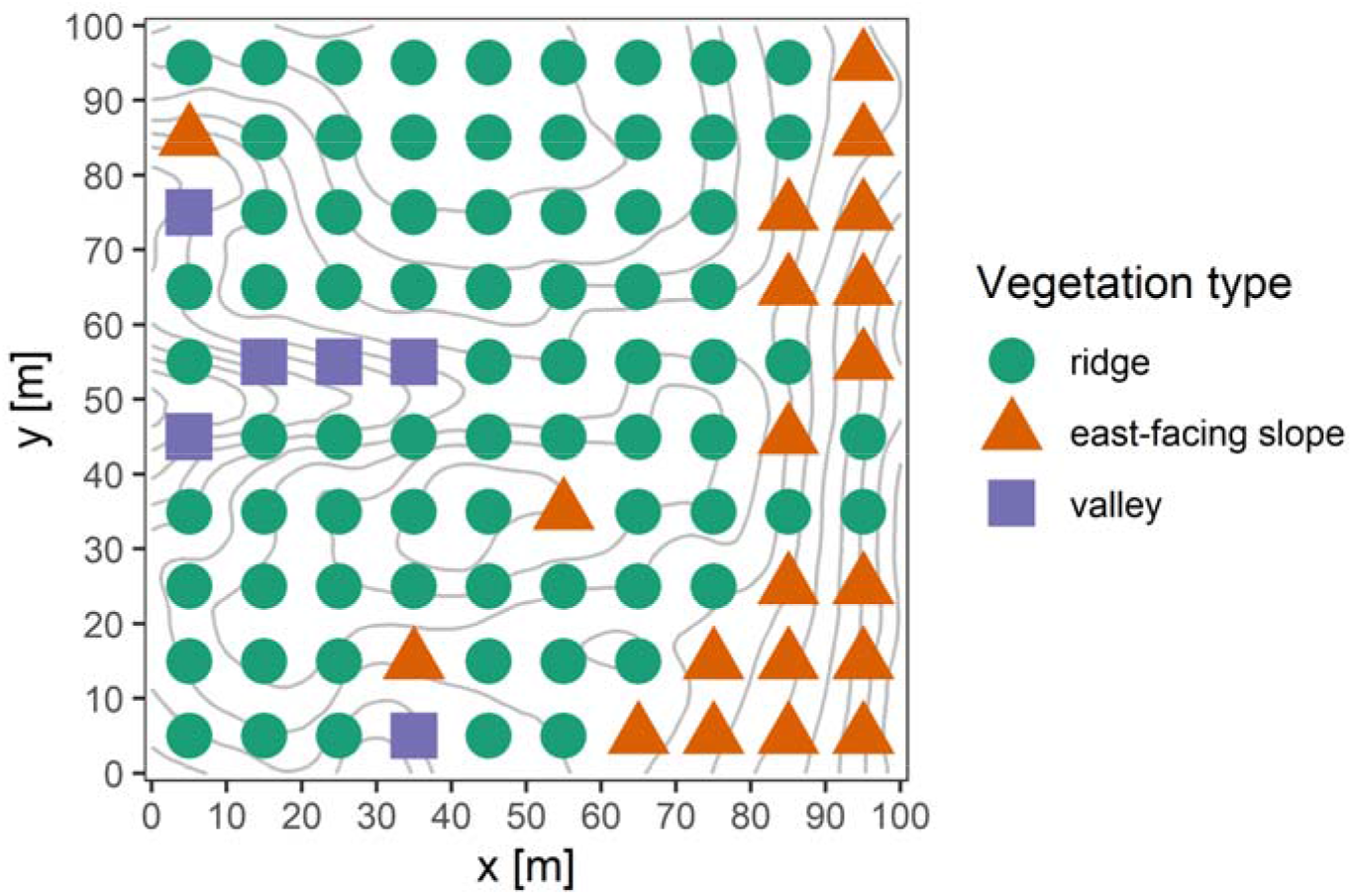
Distribution of the three vegetation types at the subplot level in LFDP.

The three vegetation types, distinguished by modified TWINSPAN, were named according to their typical topographical features and by the combination of the most diagnostic and the most dominant species as following: (1) ridge type (*Daphniphyllum himalayense* subsp. *macropodum*-*Chamaecyparis obtusa* var. *formosana*), (2) east-facing slope type (*Pourthiaea villosa* var. *parvifolia*-*Rhododendron formosanum*) and (3) valley type (*Hydrangea angustipetala*-*Eurya crenatifolia*). Their compositional, physiognomic and environmental characteristics are described below.

**Ridge type** (*Daphniphyllum himalayense* subsp. *macropodum*-*Chamaecyparis obtusa* var. *formosana*) is the main vegetation type of LFDP, mostly distributed in the subplots on the wide ridge in the west and middle part of LFDP, containing 74 subplots. Diagnostic species of ridge type include *Daphniphyllum himalayense* subsp. *macropodum* and *Rhododendron formosanum* (listed by decreasing fidelity; Appendix S1: Table S1); dominant species include *Rhododendron formosanum*, *Chamaecyparis obtusa* var. *formosana*, *Quercus sessilifolia*, *Trochodendron aralioides*, *Prunus transarisanensis*, *Quercus longinux*, *Ilex tugitakayamensis*, *Cleyera japonica* and *Acer palmatum* var. *pubescens* (listed by decreasing dominance); and constant species include *Trochodendron aralioides*, *Neolitsea acuminatissima*, *Chamaecyparis obtusa* var. *formosana* and *Cleyera japonica* (listed by decreasing constancy).

**East-facing slope type** (*Pourthiaea villosa* var. *parvifolia*-*Rhododendron formosanum*) subplots mostly distribute on the east facing windward slopes, containing 20 subplots, diagnostic species of east-facing slope type include *Pourthiaea villosa* var. *parvifolia*, *Eurya glaberrima*, *Viburnum luzonicum*, *Quercus stenophylloides*, *Microtropis fokienensis*, *Osmanthus heterophyllus*, *Tetradium ruticarpum*, *Ilex sugerokii* var. *brevipedunculata*, *Itea parviflora*, *Litsea elongata* var. *mushaensis* and *Skimmia japonica* subsp. *distincte-venulosa* (Appendix S1: Table S1); dominant species include *Rhododendron formosanum*, *Quercus sessilifolia*, *Quercus longinux* and *Neolitsea acuminatissima*; and the constant species include *Symplocos macrostroma*, *Eurya crenatifolia*, *Rhododendron formosanum*, *Neolitsea acuminatissima*, *Quercus sessilifolia*, *Chamaecyparis obtusa* var. *formosana* and *Camellia brevistyla*.

**Valley type** (*Hydrangea angustipetala*-*Eurya crenatifolia* type) subplots mostly distribute on the valley slope in the west part of LFDP, containing 6 subplots, diagnostic species of valley type include *Hydrangea angustipetala* (Appendix S1: Table S1); dominant species include *Quercus sessilifolia*, *Eurya crenatifolia*, *Cleyera japonica*, *Chamaecyparis obtusa* var. *formosana*, *Camellia brevistyla* and *Acer palmatum* var. *pubescens*; and constant species include *Symplocos macrostroma*, *Eurya crenatifolia*, and *Quercus sessilifolia*.

In total, the ridge type contains 56 species and 3160 individuals, the east-facing slope type contains 51 species and 1941 individuals, and the valley type contains 25 species and 119 individuals. When considering statistics of individual 10 m × 10 m subplots, the differences between the vegetation types are as follows. BA for the ridge type is, on average, 74.1 m^2^ ha^−1^ (11.3–191.6 m^2^ ha^−1^), for the east-facing slope type 63.1 m^2^ ha^−1^ (26.8–94.8 m^2^ ha^−1^), and for the valley type 27.4 m^2^ ha^−1^ (5.5–77.2 m^2^ ha^−1^), which is significantly lower than the other two types (Fig. S1a). Mean DBH for the ridge type is, on average, 15.8 cm (6.7–29.5 cm), for east-facing slope type 9.8 cm (6.3–14.9 cm), and for the valley type 12.7 cm (6.1–26.5 cm), meaning that the ridge type is significantly higher than east-facing type, but valley type is not significantly different from the other two types (Fig. S1b). Density of individuals for ridge type is, on average, 42.7 (11–143 individuals), for east-facing slope type 97.1 (28–184 individuals), and for valley type 19.8 (14–30 individuals), with east-facing slope type being significantly higher than the other two types (Fig. S1c). Species richness of ridget type is, on average, 14 species (7–25 species), for east-facing ridget type is 21.9 species (14–31 species), and for valley type is 11.3 species (7–6 species), with east-facing slope type being significantly richer than the other two types (Fig. S1d). When considering mean BA of different leaf habit types (conifer, evergreen broadleaf and deciduous broadleaf), there are some consistent difference between vegetation types. Mean BA of coniferous species is highest in the ridge type (27.6 m^2^ ha^−1^), compared to the significantly lower values in the east-facing slope type (3.4 m^2^ ha^−1^) and not-significantly different valley type (12.0 m^2^ ha^−1^)(Fig. S1e). Mean BA of evergreen broadleaf species is highest in the east-facing slope type (56.9 m^2^ ha^−1^), which is significantly higher than ridge type (42.2 m^2^ ha^−1^), and significantly lowest values are in the valley type (13.9 m^2^ ha^−1^)(Fig. S1f). Differences in mean BA of deciduous broadleaf species between types were not significant (Fig. S1g). Considering differences in environmental variables between vegetation types (Fig. S2a–f), ridge type occurs in subplots with higher elevation and convexity, with milder slopes, weaker windwardness, and lower soil rockiness and soil pH. East-facing slope type occurs in subplots with relatively lower elevation, high convexity, steeper slopes, significantly higher windwardness, low soil rockiness and intermediate soil pH. Valley type occurs in subplots with lower elevation, negative convexity, steeper slopes, low windwardness, higher soil rockiness, and higher soil pH. There are no significant differences in soil depth between different vegetation types (Fig. S2g).

The result of DCA shows the main gradients in species composition within the plot, the relationships between vegetation types, and also the relationship of vegetation patterns to environmental factors (passively projected onto ordination diagram, Fig. 5). The main gradient in species composition (the first DCA axis) is negatively related to windwardness and positively to elevation, and separates north-east facing vegetation types on windward slopes from the ridge vegetation type in relatively higher elevation within the plot. This pattern can be clearly observed when the site scores of individual subplots are visualised as bubble diagram in the area of the plot (Fig. 6a). The second main gradient in species composition (the second DCA axis) is negatively related to convexity and positively to soil rockiness, and separates subplots of the valley vegetation type with concave topography and high soil rockiness, from the flat or convex subplots of the other two vegetation types with lower or no soil rockiness (see also bubble diagram on Fig. 6b). The four soil variables significantly related to the first two DCA axes (pH, Cu, Mn and P) are separating valley type and partly also east-facing type with higher pH, Mn and Cu and lower P, from the ridge type with lower pH, Mn and Cu, but higher P. DBH size-class analysis for those species having at least 15 individuals (30 out of 65 species) revealed that most species have continuous regeneration and belong to J-shaped pattern (7 species, e.g. *Chamaecyparis obtusa* var. *formosana* and *Trochodendron aralioides*) or L-shaped pattern (12 species, e.g. *Neolitsea acuminatissima*, *Rhododendron formosanum*, and *Symplocos macrostroma*). In contrast, some species have fluctuating patterns (11 species, e.g. *Daphniphyllum himalayense* subsp. *macropodum*, *Dendropanax dentiger*, and *Prunus transarisanensis*), indicating periodic regeneration. Distribution maps and histograms of DBH size-class patterns for each species are at https://github.com/zdealveindy/VegLab/tree/main/data/lalashan-fdp-vegetation.

**Figure 5.**
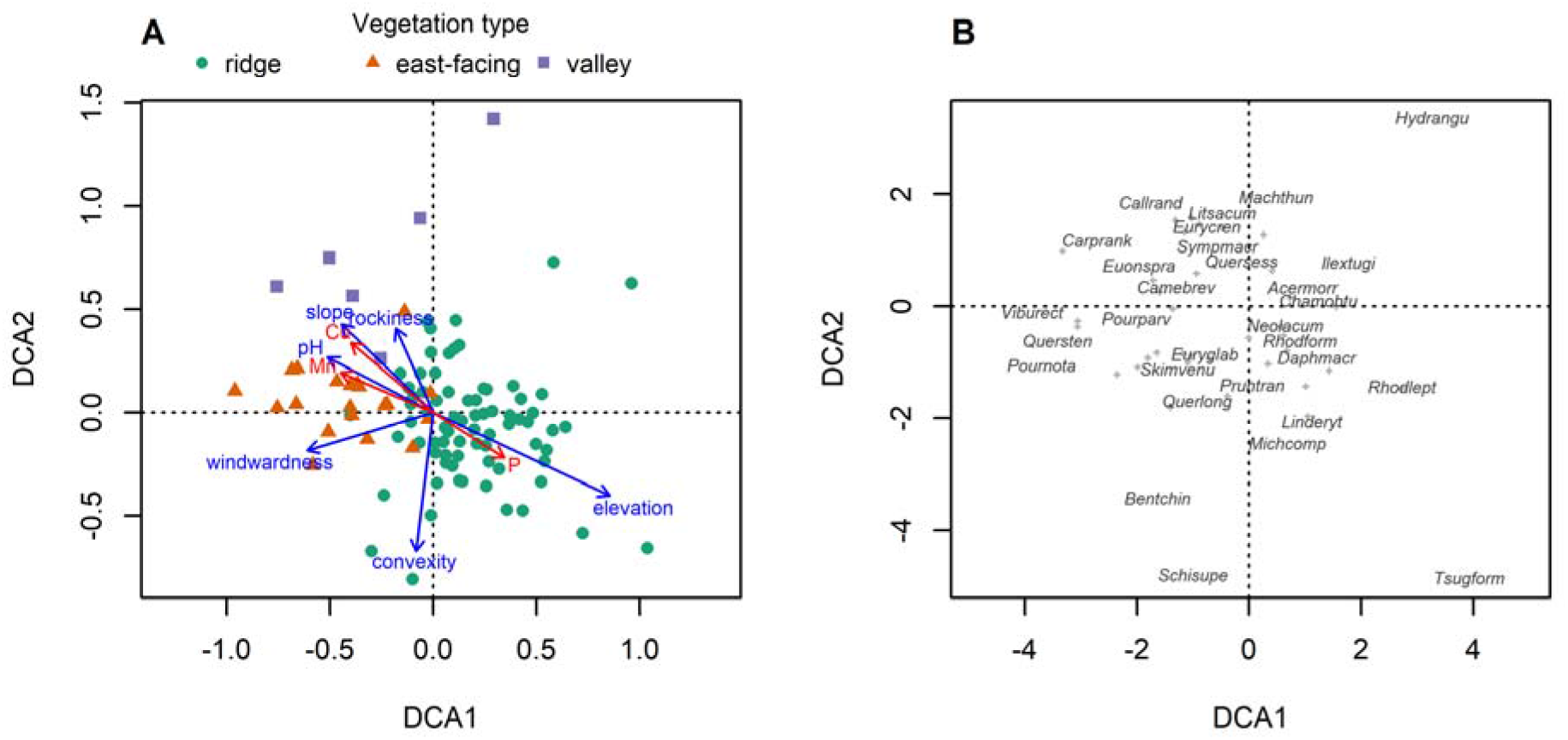
DCA ordination diagrams, showing (A) the relationships of the three vegetation types and environmental factors, passively projected onto the DCA ordination (blue vectors for variables measured in 100 subplots and significant at P < 0.05, red vectors for variables measured in 25 subplots and significant at P < 0.1), and (B) scores of the more dominant species (for the meaning of abbreviations, see Supplement, Table S3).

**Figure 6.**
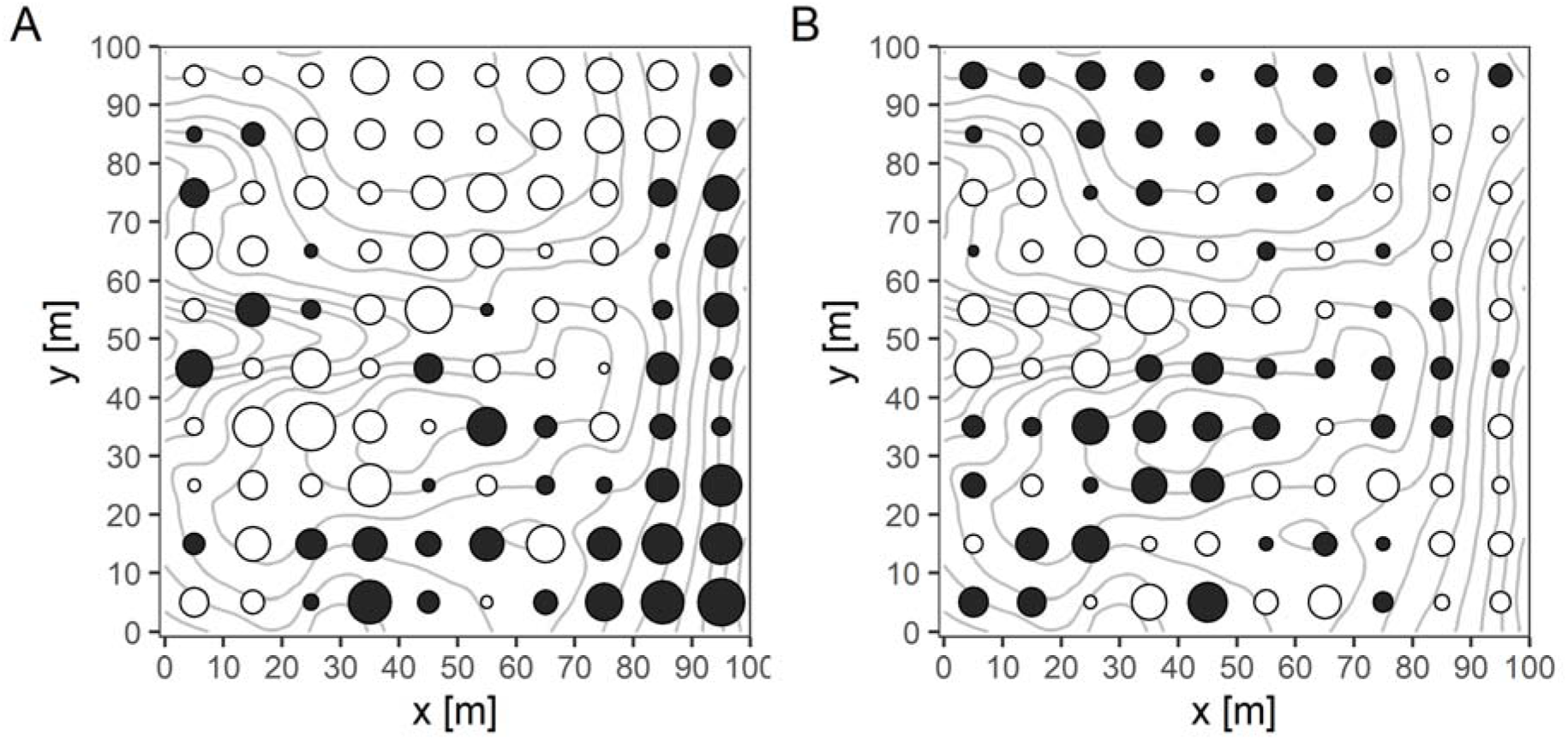
Values of site scores along the first (A) and second (B) DCA axis, displayed in the space of the forest dynamics plot. The size of the bubble indicates absolute value of the score, the colour indicates negative (black) and positive (white) value.

Regarding the multiple regression on *z*-score standardised teabag decomposition data, decomposition rate *k* can be best explained negatively by the concentration of Fe in the soil and positively by the total BA of evergreen tree species in each subplot (*k =* −0.547 × Fe *+* 0.376 × BA_evergreen_, R^2^ = 0.366, P = 0.007; Fig. S4 A&B). The stabilisation index *S* is positively related to northeasterness and to the total BA of coniferous species in each subplot (*S =* 0.757 × northeasterness + 0.588 × BA_coniferous_, R^2^ = 0.571, P < 0.001, Fig. S4 C&D).

The mean annual precipitation, measured by Lalashan saddle weather station between January 1 to December 31, 2021, was 4024 mm/year. The mean annual temperature was 13.7 °C, the mean monthly temperature of the warmest month (July) was 18.9 °C, and the mean monthly temperature of the coldest month (January) was 4.8 °C (Fig. 7). Frost events occurred in January, February and December. The prevailing wind direction measured between January 1 and December 31, 2021, was mainly from northeast (around 60°) and occasionally also from southwest (around 240°; Fig. 8). The number of foggy days (i.e. days with at least one measurement of visibility below 1000 m) was 335 days in average per year from January 2021 to December 2021. The total duration time of fog was 35.0% time of year. In winter, the fog occurred frequently and the duration time was higher than 45% time of each month (Fig. 9).

**Figure 7.**
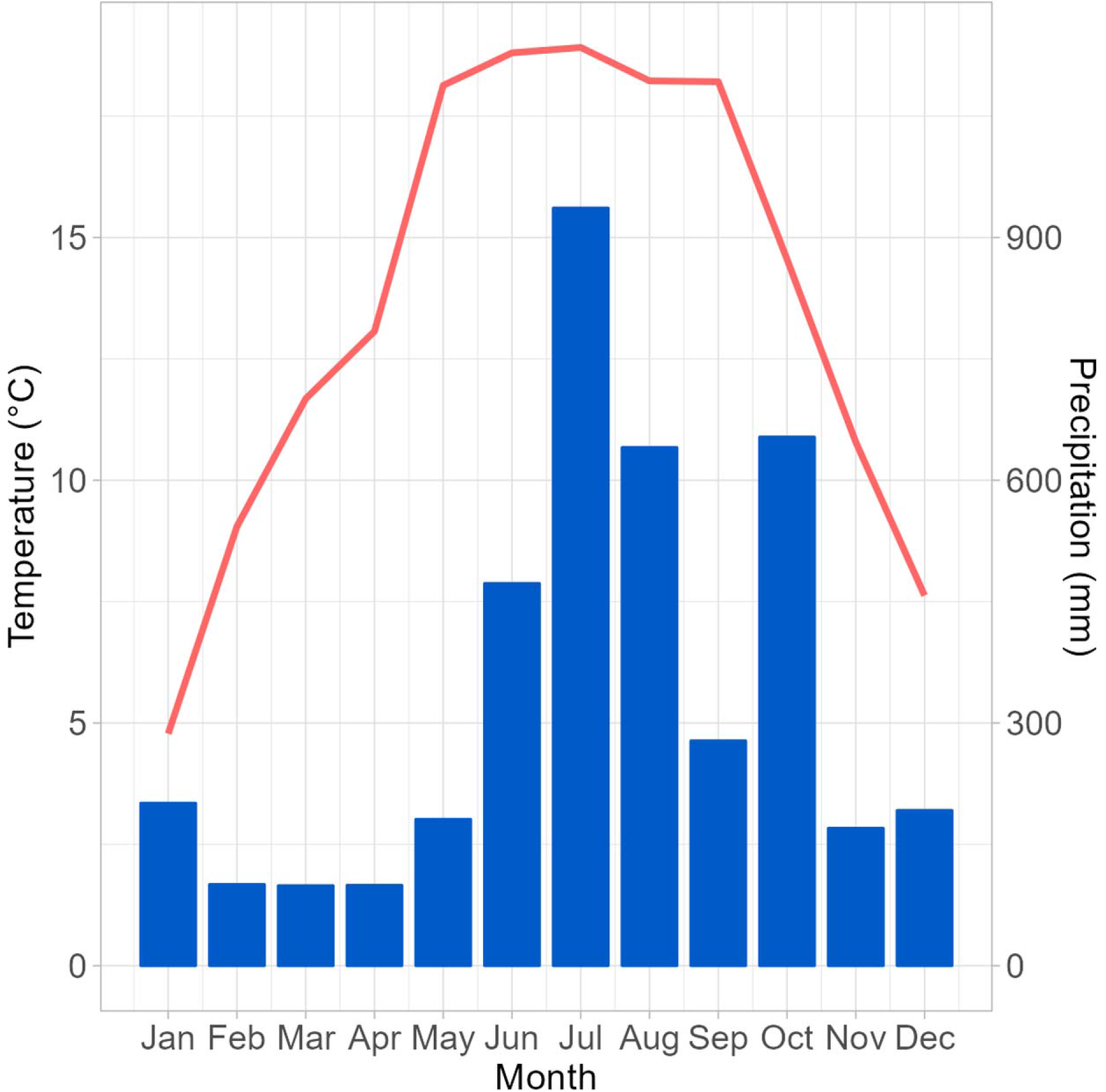
Monthly temperatures (red curve, scale on the left) and precipitations (blue bars, scale on the right), measured by Lalashan Saddle weather station between January 2021 and December 2021.

**Figure 8.**
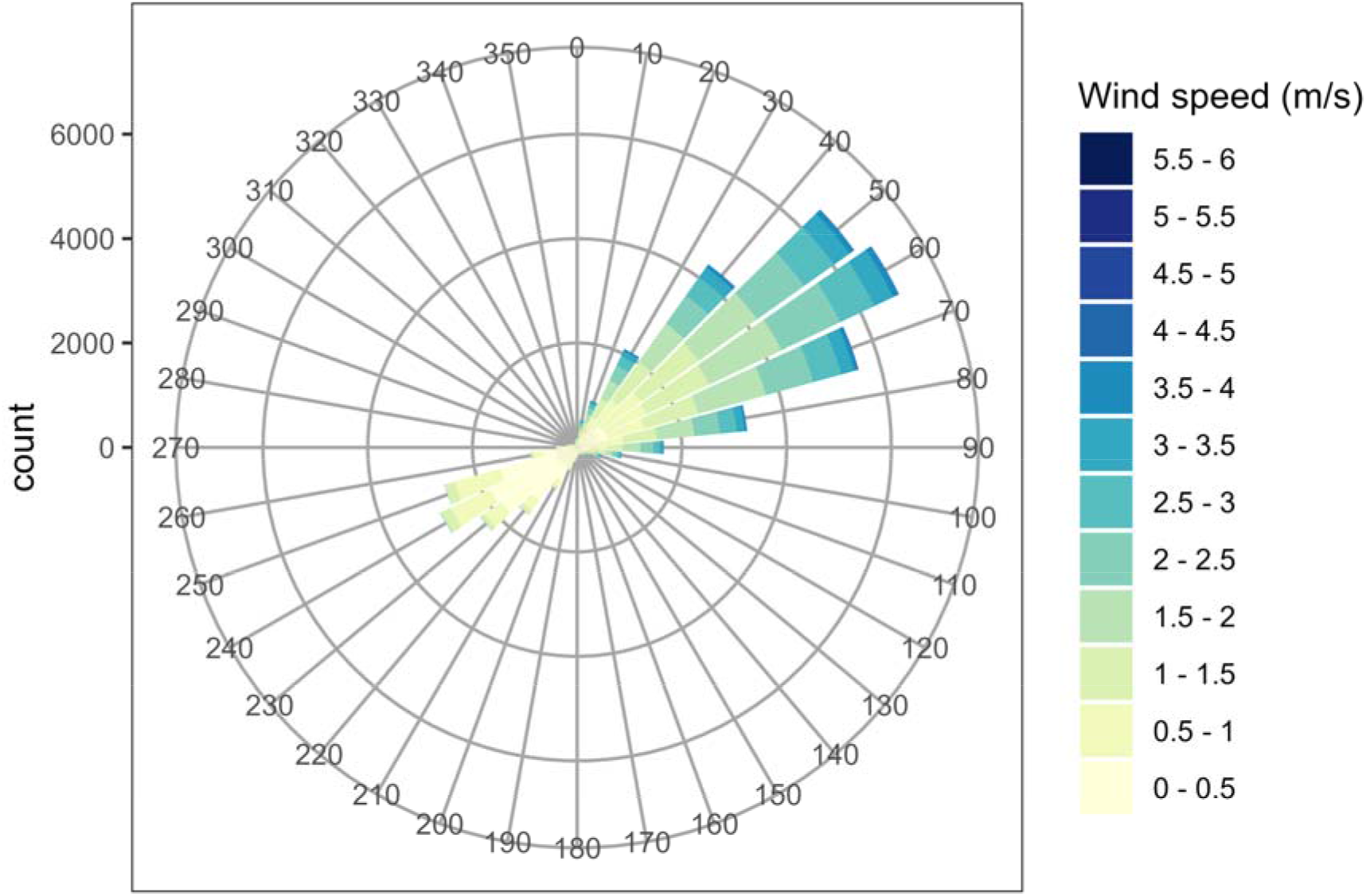
Wind rose charts from Lalashan Saddle weather station, measured between January 1, 2021, and December 31, 2021.

**Figure 9.**
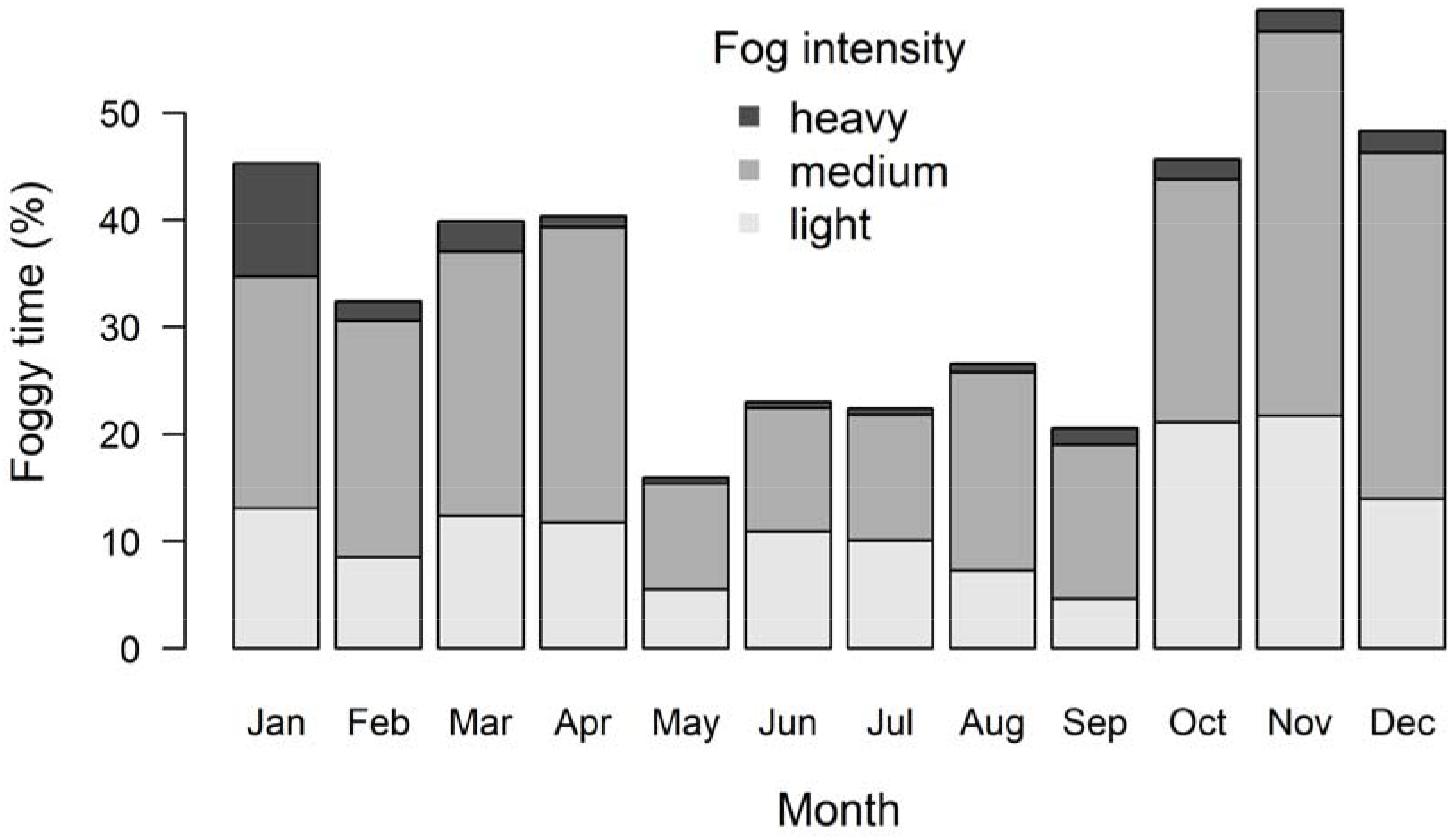
Percentages of different degrees of fogginess, as measured by visibility logger between January 1, 2021, and December 31, 2021. Light fogginess represents visibility from 500 to 1000 m, medium fogginess from 100 to 500 m, and heavy fogginess from 0 to 100 m.

For microclimate measured inside the LFDP between August 2021 and July 2023, the warmest month was July in 2022 with mean temperature 19.2°C, and the coldest month was December in 2022 with mean temperature 6.5°C (Fig. S5 A). The daily temperature deviation between valley and flat ridge, and between east-facing slope and flat ridge was generally lower than zero across a year except November and February (Fig. S5 B). Interestingly, during December and January, east-facing slopes affected by NE monsoon were slightly colder than both flat ridge and the valley, while in the other parts of the year both valley and east-facing slopes were colder than flat ridge, with valley colder than east-facing slope. The proportions of wet days from November to February were higher than 80% except the flat ridge in February (Fig. S6 A), and dry days were not frequent (Fig. S6 B). The soil moisture between April 20, 2022 and May 20, 2022 increased along the topographical gradient from the convex ridge to concave valley (Fig. S7).

## Discussion

Analysis of our data shows that the variation in composition of woody species in LFDP is driven by two main environmental gradients, windwardness and convexity. Change of vegetation along windwardness gradient happens on compositional but also diversity and physiognomic levels. Wind speed and direction measurement from the weather station beside the plot shows that the prevailing wind in the area is from NE (50-60°), mainly because this is the overall direction of the winter monsoon. Windward east-facing slope vegetation type has the highest number of diagnostic species preferring this wind-affected habitat, has subplots with the highest richness and also with highest stem and branch density, creating dense but short canopy forest, which is being avoided by coniferous species but somewhat preferred by deciduous ones. It is commonly observed that forest becomes shorter and denser under the influence of chronic wind (Lawton 1982), and similar dense stands can also be found in other windward-type forests in Taiwan affected by northeast monsoon, including the forest dynamics plots at Mt. Lopei (Lin et al. 2005) and Lanjenchi (Chao et al. 2007, 2010; Ku et al. 2021). For completeness, we also have to acknowledge that the east-facing slope would be colder and wetter in general since it receives a relatively lower amount of solar radiation compared to parts of the plots facing south or southwest aspects. Also, the east-facing habitat includes steepest quadrats in the whole dynamic plot, and the steepness itself can affect forest composition and physiognomy (Moeslund et al. 2013). The ridge vegetation type, which grows in less wind-affected and more convex habitats, has on the other side tall canopy, with dominance of coniferous *Chamaecyparis obtusa* var. *formosana*, and hosts trees with the largest DBH in the plot. The valley type occurs in a relatively small area, is concentrated into convex habitats with rather steep slopes of dry gullies, and is characterised by low basal area, density and species richness of trees.

Temperature measurements from loggers in each of the three vegetation types showed that both east-facing and valley types when compared to the ridge type, are relatively colder, mainly during the spring and summer months, and partly also during winter. This pattern probably reflects an interaction between the microclimate and topography of the three habitat types. Windward east-facing habitats are colder partly because of the cooling effect of wind (mostly during the winter and spring during the winter monsoon period), but also because they receive solar radiation mostly during the morning hours, less so in the afternoon when they are already shaded. The valley type in a more topographically shaded position is generally less irradiated during the day and may also be affected by cold airflow at night. Ecological difference between convex and concave habitats is also strongly linked to soil moisture, as shown by our soil moisture measurements. The convex sites in the central part of the plot are filled with permanently wet mud (we call them mud pools on the LFDP map in Fig. 2), and during rainy periods, they even get flooded.

Although soil variables are relatively poorly linked to species composition, a few interesting patterns exist. Soil pH is consistently lowest in ridge vegetation type, probably for two reasons. First is the dominance of conifers in this habitat and, consequently, the accumulation of hard-to-decompose coniferous litter, leading to lower pH and higher C/N ratio in these subplots (Finzi et al. 1998; Satti et al. 2003; Hobbie et al. 2006). The ridge type is also mostly on elevated flat ridges prone to leaching, removing cations from the soil and reducing soil pH. Interestingly, in the ordination diagram (Fig. 5), phosphorus is negatively related to soil pH, which goes against the typical pattern where P is less available in soils with lower pH due to high levels of aluminium and iron cations, which combine with P and restrict its solubility (SanClements 2010). We speculate that the negative correlation of P and pH in our plot results from the slow litter decomposition and accumulation of P in undecomposed organic material in a form unavailable for plants, especially in ridge habitat convex subplots in higher elevation dominated by conifers. Unpublished data from our small-scale application of resin soil probes (Plant Root Simulator, Western Ag. Innovations Inc., Saskatoon, Saskatchewan, Canada) in the plot during the 2022 season supports this interpretation. It showed that the supply rate of P anions in the soil in the six buried sets of probes is generally below the method’s detection limit, indicating a strong P limitation. Finally, steeper subplots of east-facing and valley vegetation types tend to have higher concentrations of Cu and Mn for reasons which remain unclear to us.

The low importance of soil properties to changes in species composition may be linked to several limitations of these measurements. Soil properties were measured only in 25 out of 100 subplots, and the reduced sample size also reduces the analytical power. However, since on the scale of our observation, soil variables tend to be highly spatially autocorrelated, even higher number of measurements may not improve the situation since the analysis would need to be adjusted for spatial autocorrelation. Also, we do not expect that overall variation in soil conditions between subplots within a rather small area of the 1-ha plot is high enough to have a marked effect on vegetation, even though the plot’s topography is relatively heterogeneous. And finally, the method of analysing the soil nutrients itself does not really reflect their real availability for plants, but merely their concentration in the soil profile extracted by relevant chemical agents in the lab (Wilcke et al. 2002; Axmanová et al. 2011). Alternative methods may alleviate this problem, such as resin probes mentioned above or more elaborated ways of collecting and analysing the soil samples.

Although the decomposition rate and stabilisation factor, as determined by the teabag decomposition experiment in our plot, were not related to species composition changes in our analysis, they were related to a combination of soil, topographical and biotic variables. A stronger effect is expressed by the stabilisation factor, which is positively related to northeasterness and BA of coniferous species in the plot. Stabilisation reflects environmental and biotic effects on converting part of the litter into recalcitrant form, not available for further decomposition (Keuskamp et al. 2013), and can result in the accumulation of soil organic matter in the soil (Wiesmeier et al. 2019). More detailed analysis of litter decomposition data is necessary, however, to better understand the complex effect of environment and vegetation on decomposition.

## Conclusions

Our study brings data about the fine-scale composition and spatial distribution of woody species in the 1-ha Lalashan Forest Dynamics Plot, an example of an old-growth stand belonging to the submontane *Chamaecyparis* mixed cloud forest. Our topographical, soil and microclimatic data show that the species composition here is mainly driven by the effect of wind and by topographical convexity, both having profound effects on species composition, species richness and forest structure of the vegetation as well as soil and microclimatic conditions. The weather station we maintain less than 100 m from the plot provides additional climatic data showing prevailing wind coming from northeast direction and frequent fog events, most common during the winter months. The compositional and environmental data will serve as a reference for future resurveys to improve our understanding of the long-term dynamics and changes in montane cloud forest vegetation. This is important, especially in the context of ongoing climate change effects that are predicted to have led to the uplifting of the cloud layer, potentially leading to significant environmental and compositional changes to cloud forest vegetation restricted to climatically highly specialised habitat conditions.

## Authors contributions

The idea was proposed by DZ and Ching-Feng Li (Woody). TC, YNL together with DZ and other members of the Vegetation Ecology Lab and volunteers established and surveyd LFDP. TC, YNL, and KSW maintained compositional and environmental data. YNL, TC and lab members from the Vegetation Ecology Lab and Soil Survey and Remediation Lab (of Prof. Zeng-Yi Hsu) analysed the soil samples. TC conducted the analyses of woody species composition and environmental factors data for the results, prepared the interpretation of results and discussion, and wrote the first draft of the manuscript as her Master thesis. PYL conducted the tea bag decomposition experiment, analysed microclimatic data and drew figures related to climate. DZ reanalysed and extended the manuscript, redraw the figures, and prepared the final version of the text. All co-authors commented on the final version of the manuscript.

## Acknowledgements

We would like to thank to all members of Vegetation Ecology Lab and other volunteers who participated in field survey. We also greatly appreciate help from workers in Lalashan Recreation Area for help and positive attitude toward our research expeditions. Thanks also go to Taiwanese Forestry Bureau for renting us the abandoned weather station in the Lalashan saddle. The research was financially supported by Taiwanese Ministry of Science and Technology (106-2621-B-002-003-MY3 and 109-2621-B-002-002-MY3).

