## Supplement for "Vegetation of *Chamaecyparis* montane cloud forest in Lalashan Forest Dynamics Plot"

The following supplementary materials are available for this article: Chen, T., Lee, Y.-N., Lin, P.-Y., Wu, K.-S. and Zelený, D. Submitted. Vegetation of *Chamaecyparis* montane cloud forest in Lalashan Forest Dynamics Plot. Taiwania

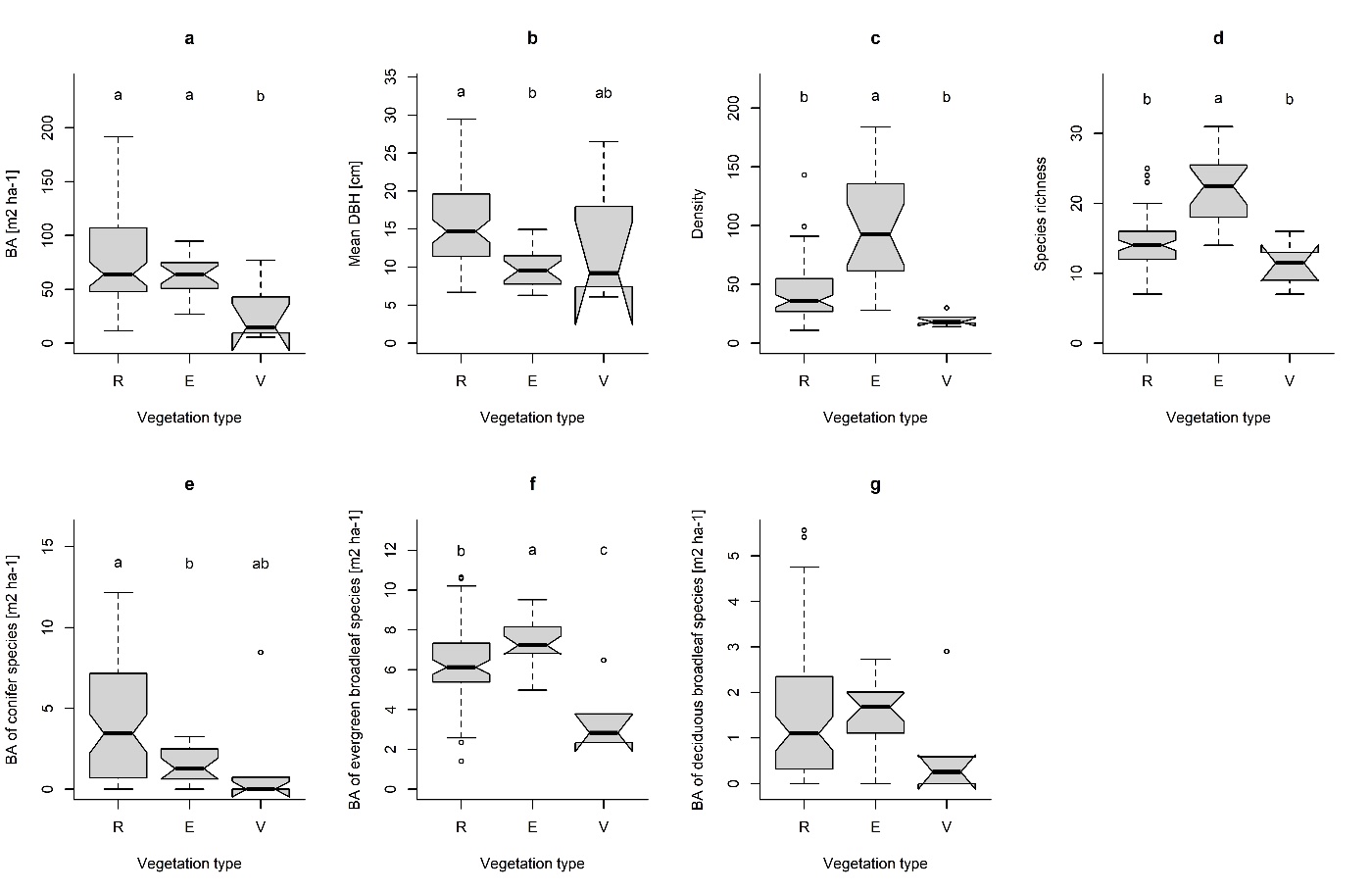

**Figure S1.** Boxplots showing the subplot-based physiognomic differences between the three vegetation types; a. BA, b. mean DBH, c. density, d. species richness, e. BA of conifer species, f. BA of evergreen broadleaf species, g. BA of deciduous broadleaf species. All differences between vegetation types, except for g, are significant (P < 0.05) through the ANOVA test. R: ridge type, E: east-facing slope type, V: valley type.

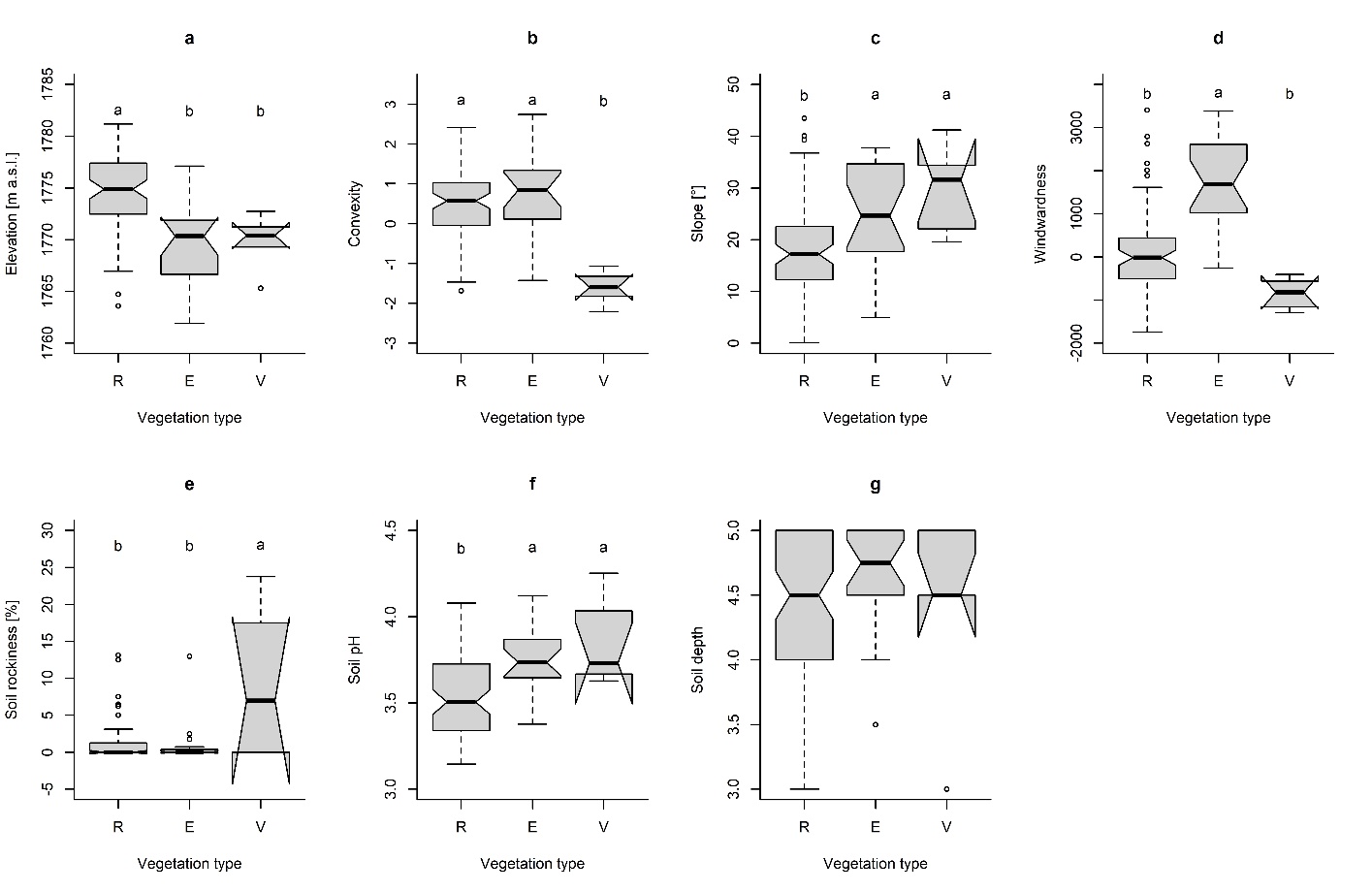

**Figure S2.** Boxplots showing the differences of different environmental factors (containing 100 subplots data) between the three vegetation types; a. elevation, b. convexity, c. slope, d. windwardness, e. soil rockiness, f. soil pH, g. soil depth. All differences between the vegetation types, except for g, are significant through ANOVA test. R: ridge type, E: east-facing slope type, V: valley type.

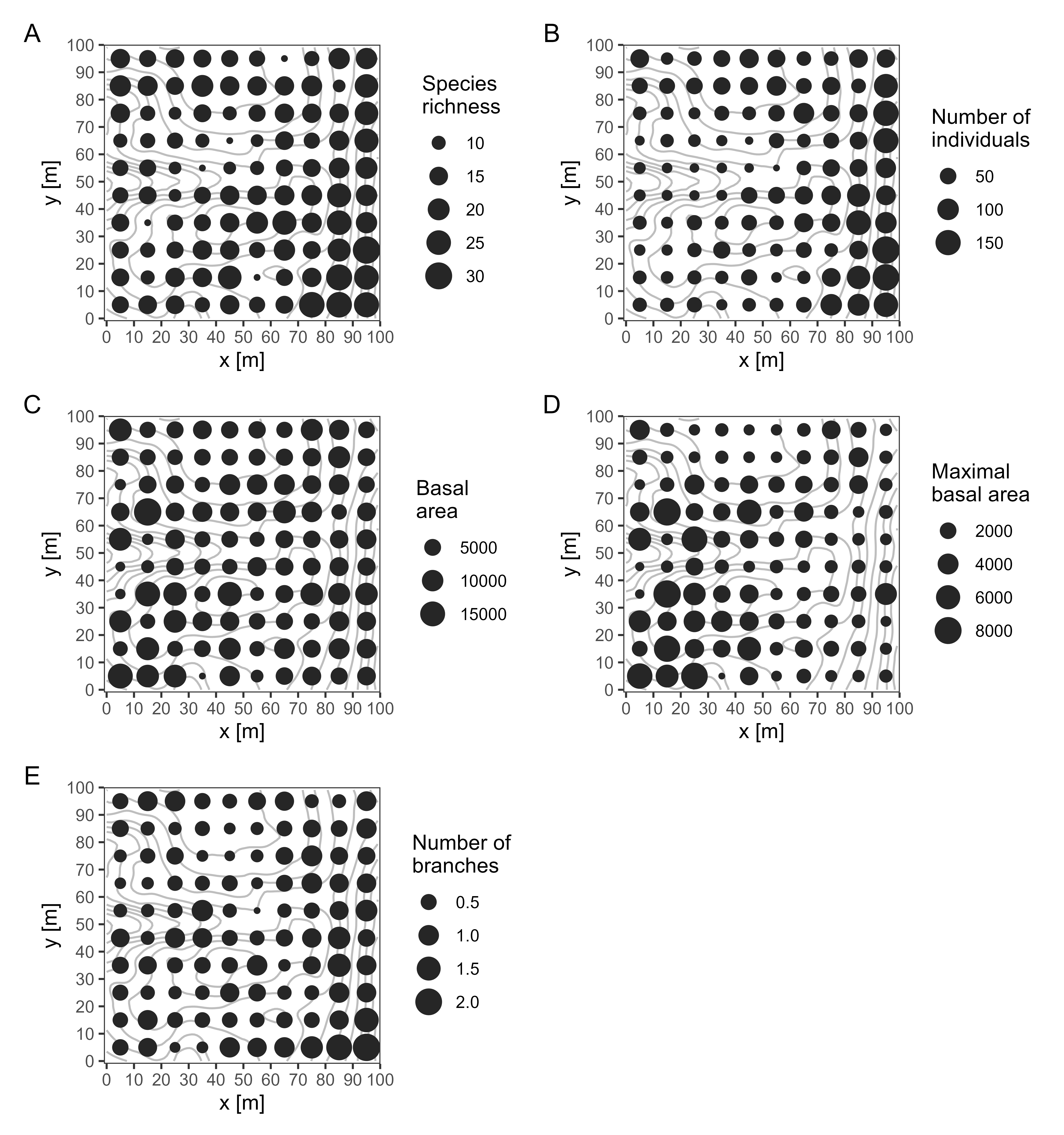

**Figure S3**. Spatial distribution of species richness (A), number of individuals per subplot (B), basal area of all individuals within the subplot (C), maximal basal area of the largest individual within the subplot (D) and average number of branches per individual within subplot (E). The size of the circle is proportional to value of the variable in given subplot.

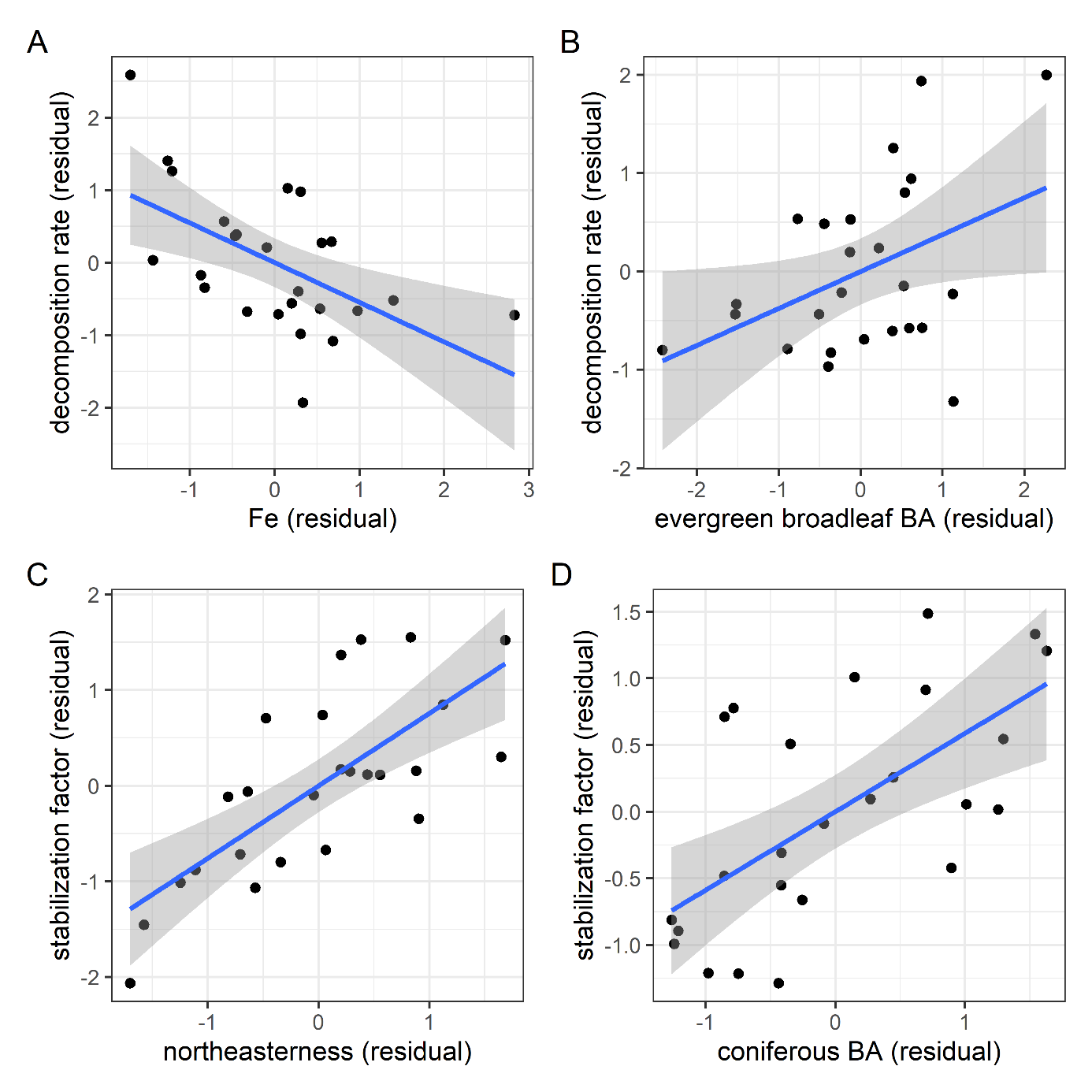

**Figure S4.** Partial regression diagrams of soil decomposition rate (A and B) and stabilisation factor (C and D) to selected significant environmental factors and biotic variables.

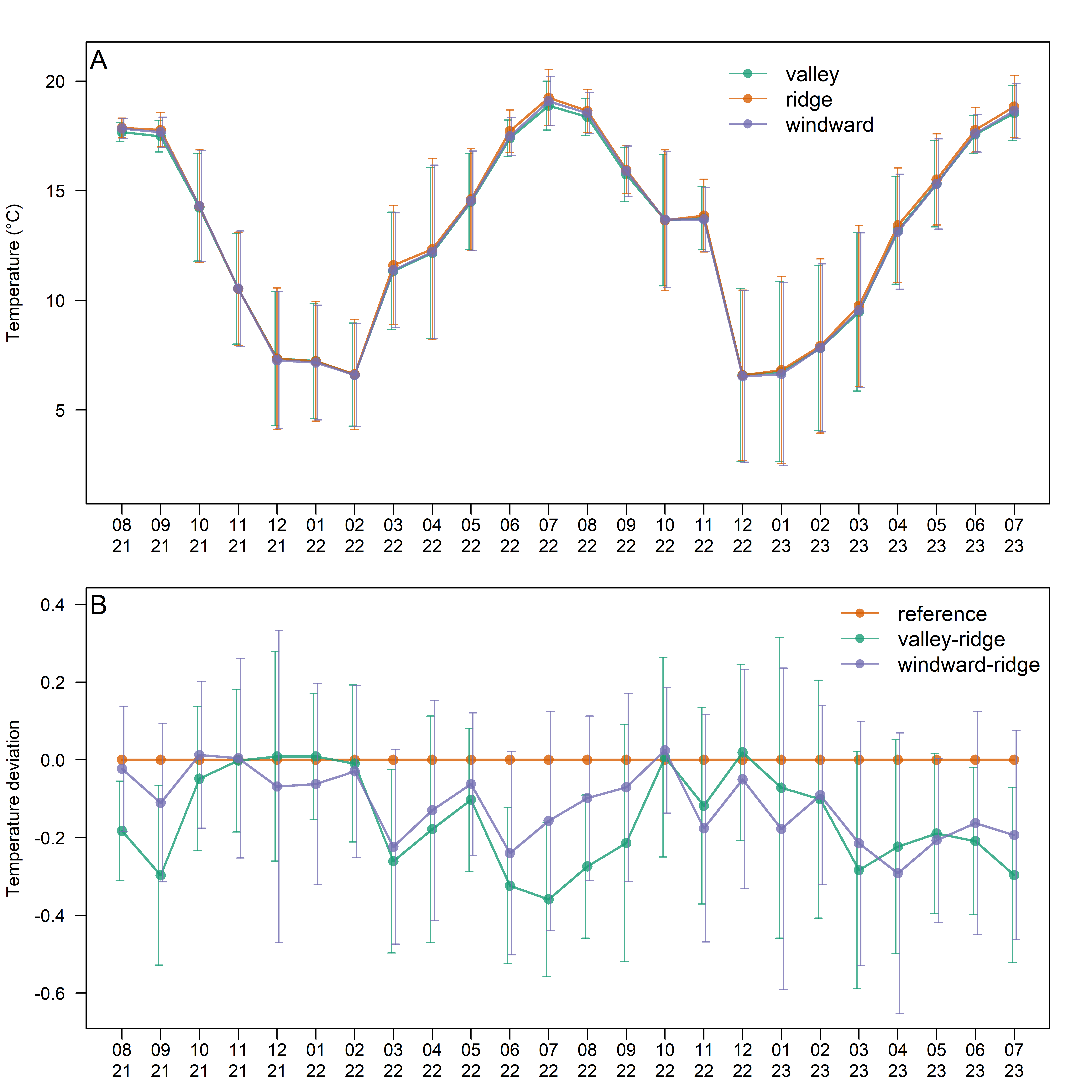

**Figure S5.** The pattern of mean monthly temperature (A) and mean monthly daily temperature deviation (B) in the LFDP from temperature measurements between August 1, 2021 and July 31, 2023. Valley: valley in subplot (0,4); ridge: flat ridge in subplot (3,2); windward: east-facing wind-affected slope in subplot (9,3).

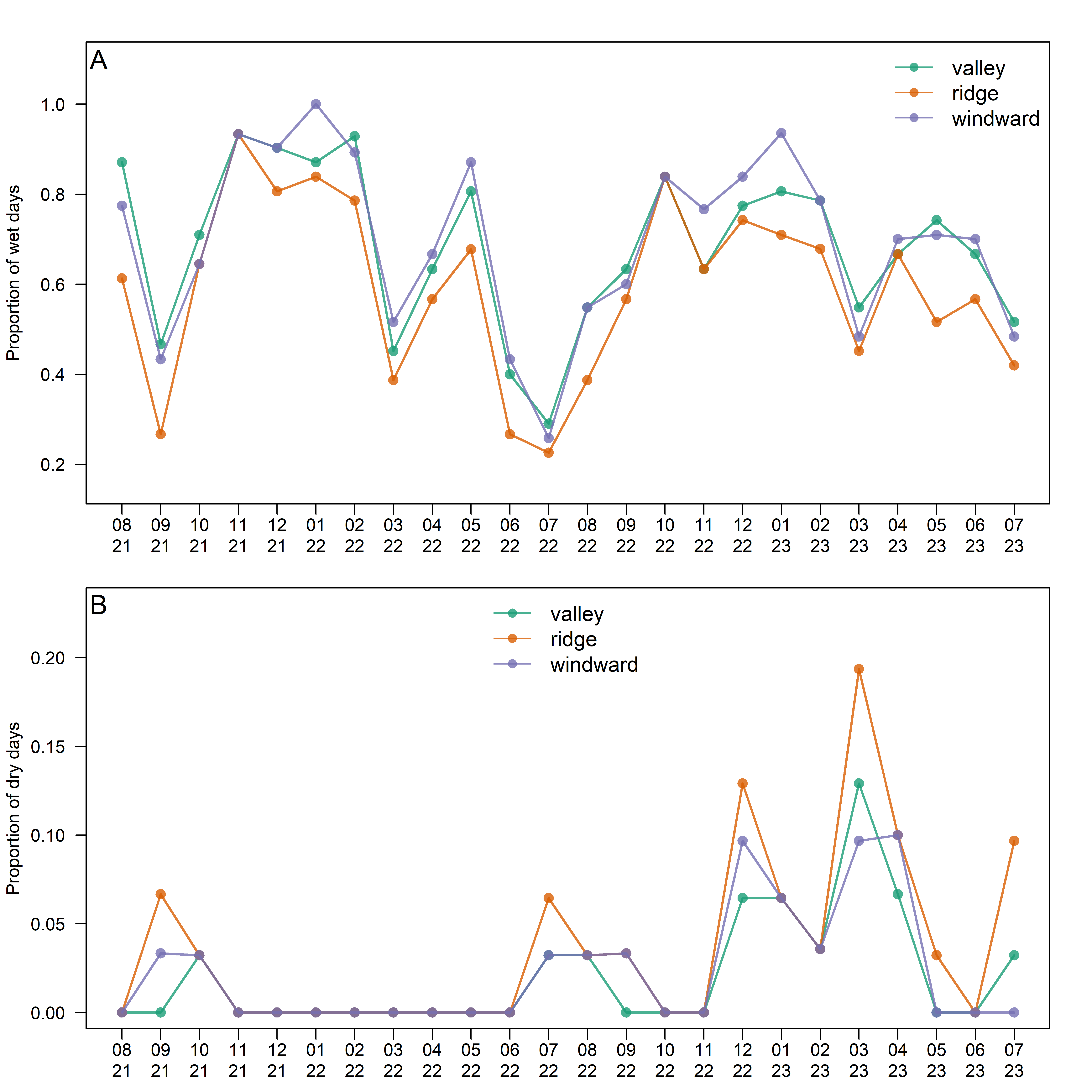

**Figure S6.** Monthly proportions of wet days (A) and dry days (B) from August 1, 2021 to July 31, 2023. Wet days are defined as days with mean RH > 99%, while dry days with mean RH < 75%. Valley: valley in subplot (0,4); ridge: flat ridge in subplot (3,2); windward: east-facing wind-affected slope in subplot (9,3).

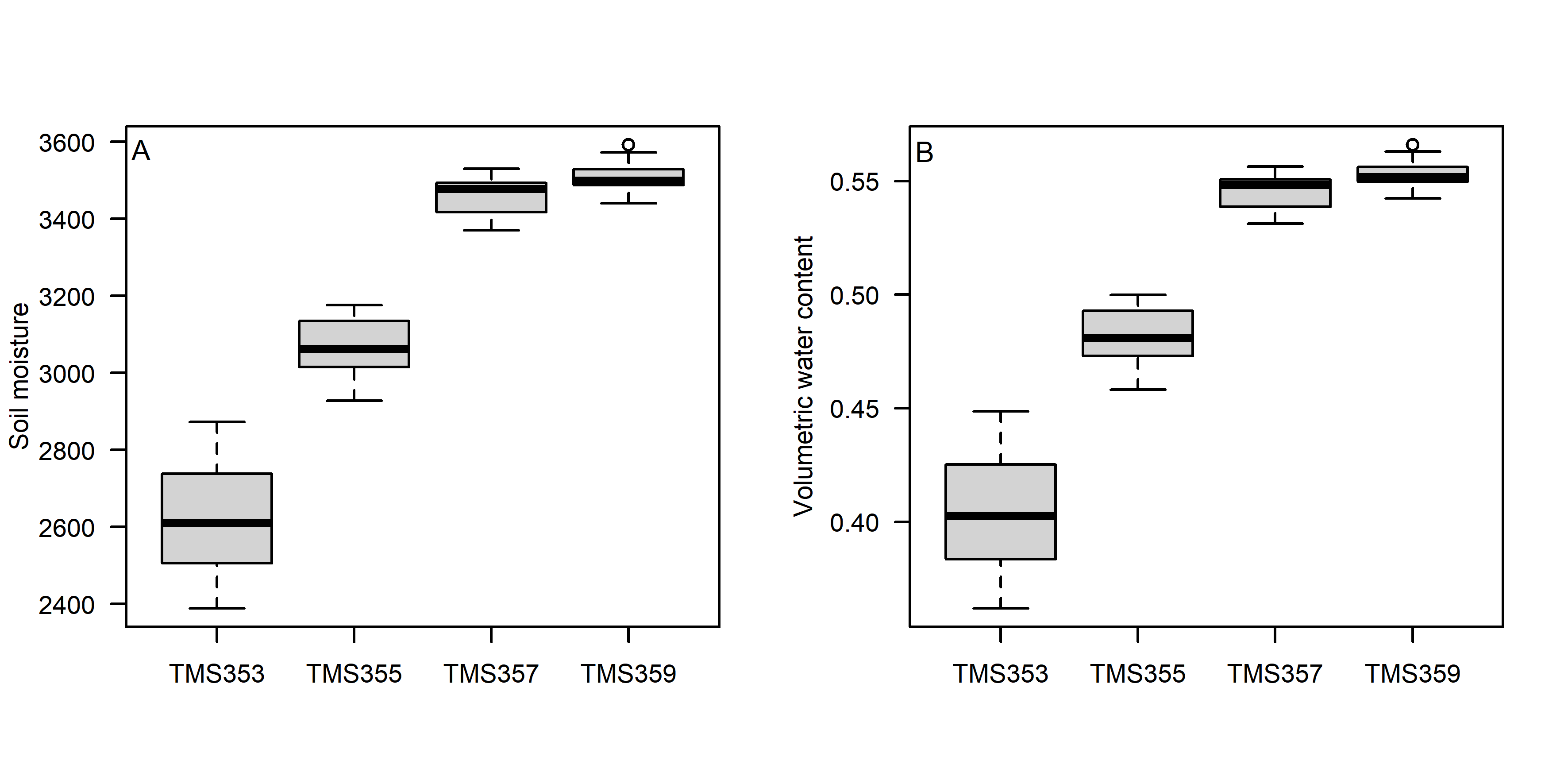

**Figure S7.** Differences in soil moisture (A) and volumetric water content (B) along the topographical gradient from the convex ridge (TMS353) to the concave valley (TMS359), measured by TOMST TMS-4 loggers close to the centre of the plot from April 17, 2022, till May 28, 2022.

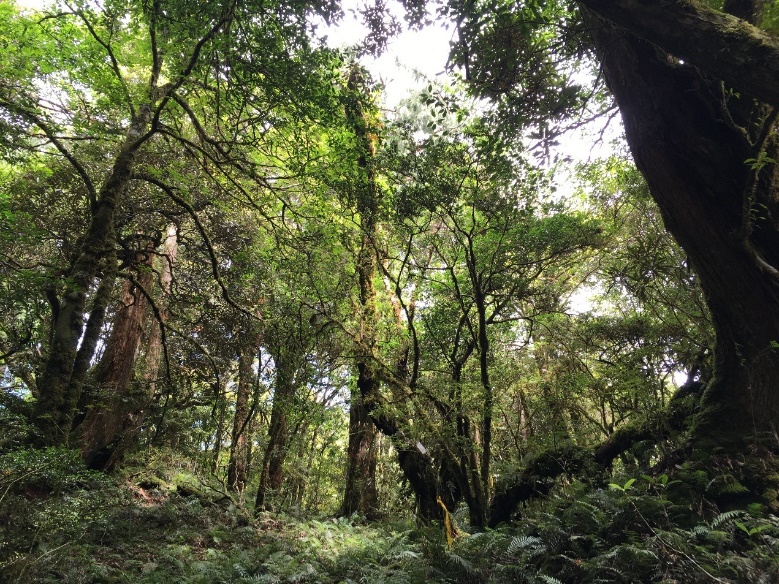

B
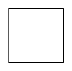

A

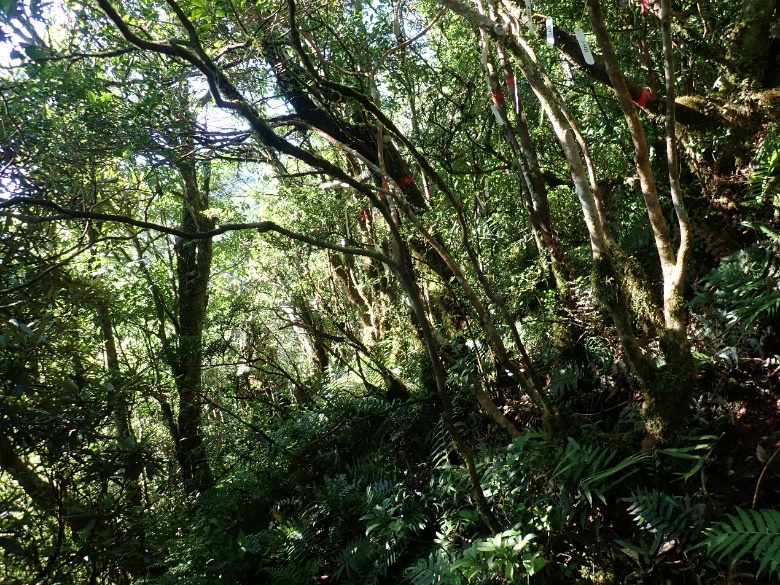

C
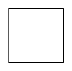

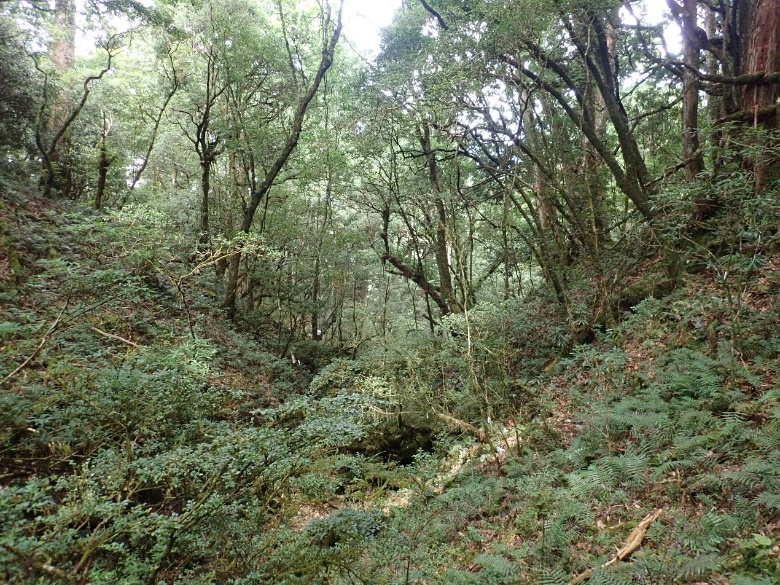

**Figure S8.** Photographs of the three vegetation types in LFDP; A - ridge type, B - east-facing slope type, C - valley type. Photo credit by Ting Chen.

**Table S1.** Diagnostic species of the three vegetation types. Values are the relative percentage frequency (Freq.), and species are sorted by decreasing fidelity (Φ). The green colour indicates diagnostic species with a fidelity ≥ 35%.

| **Vegetation type** | **R** | |  | **E** | |  | **V** | |
| --- | --- | --- | --- | --- | --- | --- | --- | --- |
| **No. of plots** | 74 | |  | 20 | |  | 6 | |
|  | Freq. | Φ. |  | Freq. | Φ |  | Freq. | Φ |
| *Daphniphyllum himalayense* subsp. *macropodum* | 62 | 38.1 |  | 30 | 33 |  | 17 | - |
| *Rhododendron formosanum* | 93 | 37.9 |  | 95 | - |  | 17 | - |
| *Pourthiaea villosa* var. *parvifolia* | 7 | - |  | 75 | 75.8 |  |  | - |
| *Eurya glaberrima* | 61 | - |  | 95 | 60.9 |  |  | - |
| *Viburnum luzonicum* | 11 | - |  | 50 | 52.3 |  |  | - |
| *Quercus stenophylloides* |  | - |  | 35 | 51.4 |  |  | - |
| *Microtropis fokienensis* | 12 | - |  | 50 | 51.1 |  |  | - |
| *Osmanthus heterophyllus* | 32 | - |  | 70 | 43.8 |  | 17 | - |
| *Tetradium ruticarpum* |  | - |  | 25 | 42.6 |  |  | - |
| *Ilex sugerokii* var. *brevipedunculata* | 4 | - |  | 30 | 41.6 |  |  | - |
| *Itea parviflora* | 3 | - |  | 25 | 38.5 |  |  | - |
| *Litsea elongata* var. *mushaensis* | 38 | - |  | 90 | 37.3 |  | 67 | - |
| *Skimmia japonica* subsp. *distincte-venulosa* | 4 | - |  | 25 | 36.6 |  |  | - |
| *Hydrangea angustipetala* | 3 | - |  |  | - |  | 33 | 46.4 |
| *Eurya loquaiana* | 7 | - |  | 10 | - |  |  | - |
| *Chamaecyparis obtusa* var. *formosana* | 85 | - |  | 90 | - |  | 33 | - |
| *Tsuga chinensis* var. *formosana* | 3 | - |  |  | - |  |  | - |
| *Neolitsea acuminatissima* | 88 | - |  | 95 | - |  | 67 | - |
| *Trochodendron aralioides* | 95 | 34.2 |  | 75 | - |  | 50 | - |
| *Cleyera japonica* | 81 | - |  | 75 | - |  | 67 | - |
| *Ilex lonicerifolia* | 12 | - |  | 30 | 20.5 |  | 17 | - |
| *Pourthiaea beauverdiana* var. *notabilis* |  | - |  | 5 | - |  |  | - |
| *Euonymus spraguei* |  | - |  | 5 | - |  |  | - |
| *Dendropanax dentiger* | 62 | - |  | 70 | - |  | 50 | - |
| *Prunus transarisanensis* | 38 | - |  | 55 | - |  |  | - |
| *Schima superba* | 1 | - |  |  | - |  |  | - |
| *Acer morrisonense* | 1 | - |  |  | - |  |  | - |
| *Michelia compressa* | 3 | - |  | 5 | - |  |  | - |
| *Litsea acuminata* | 8 | - |  | 5 | - |  | 17 | - |
| *Lindera erythrocarpa* | 11 | - |  | 10 | - |  |  | - |
| *Photinia niitakayamensis* | 8 | - |  |  | - |  |  | - |
| *Acer kawakamii* | 8 | - |  | 10 | - |  |  | - |
| *Quercus sessilifolia* | 72 | - |  | 90 | - |  | 83 | - |
| *Sycopsis sinensis* | 5 | - |  | 5 | - |  | 17 | - |
| *Rhamnus crenata* | 8 | - |  | 5 | - |  |  | - |
| *Neolitsea aciculata* | 9 | - |  | 5 | - |  |  | - |
| *Styrax formosanus* | 14 | - |  | 10 | - |  |  | - |
| *Machilus thunbergii* | 3 | - |  |  | - |  |  | - |
| *Acer palmatum* var. *pubescens* | 35 | - |  | 50 | - |  | 33 | - |
| *Ilex tugitakayamensis* | 30 | - |  | 35 | - |  |  | - |
| *Symplocos macrostroma* | 72 | - |  | 100 | 22.9 |  | 100 | - |
| *Carpinus rankanensis* | 4 | - |  | 30 | 24.7 |  | 17 | - |
| *Symplocos formosana* | 16 | - |  | 60 | 25.7 |  | 50 | - |
| *Ligustrum liukiuense* | 53 | - |  | 80 | 20.3 |  | 67 | - |
| *Eurya crenatifolia* | 73 | - |  | 100 | 22.2 |  | 100 | - |
| *Ilex suzukii* |  | - |  | 10 | 26.3 |  |  | - |
| *Viburnum foetidum* var. *rectangulatum* |  | - |  | 15 | 32.4 |  |  | - |
| *Sorbus randaiensis* | 3 | - |  | 20 | 33.2 |  |  | - |
| *Barthea barthei* | 3 | - |  | 20 | 33.2 |  |  | - |
| *Viburnum urceolatum* |  | - |  | 10 | 26.3 |  |  | - |
| *Ilex hayatana* | 31 | - |  | 55 | 30.9 |  | 17 | - |
| *Camellia brevistyla* | 39 | - |  | 85 | 31.4 |  | 67 | - |
| *Rhododendron leptosanthum* | 1 | - |  |  | - |  |  | - |
| *Tetradium glabrifolium* | 1 | - |  |  | - |  |  | - |
| *Viburnum sympodiale* | 27 | - |  | 40 | - |  | 17 | - |
| *Quercus longinux* | 26 | - |  | 45 | - |  |  | - |
| *Vaccinium bracteatum* | 1 | - |  |  | - |  |  | - |
| *Rhododendron pseudochrysanthum* | 1 | - |  |  | - |  |  | - |
| *Prunus phaeosticta* | 23 | - |  | 35 | - |  | 67 | - |
| *Berberis hayatana* |  | - |  | 5 | - |  |  | - |
| *Symplocos migoi* |  | - |  |  | - |  | 17 | - |
| *Callicarpa randaiensis* | 5 | - |  | 30 | 11.9 |  | 33 | - |
| *Pieris taiwanensis* | 1 | - |  |  | - |  |  | - |
| *Benthamidia japonica* var. *chinensis* | 1 | - |  |  | - |  |  | - |
| *Chamaecyparis formosensis* | 1 | - |  | 10 | - |  |  | - |

**Table S2.** The multiple regression between each environmental variables and the first two DCA axes. No. subplots = in how many subplots the variable was measured; DCA1 and DCA2 = directional cosines of each variable, *r*^2^ = coefficient of determination; *P*-value = significance (calculated by Monte Carlo permutation test with toroidal shift; *** = P < 0.001; ** = P < 0.01; * = P < 0.05; . = P < 0.1); No. perm. = number of permutations in the test.

| Variable | No. subplots | DCA1 | DCA2 | *r*^2^ | *P*-value |  | No. perm. |
| --- | --- | --- | --- | --- | --- | --- | --- |
| elevation | 100 | 0.904 | -0.427 | 0.436 | 0.0025 | ** | 399 |
| convexity | 100 | -0.124 | -0.992 | 0.225 | 0.0025 | ** | 399 |
| slope | 100 | -0.720 | 0.694 | 0.186 | 0.0250 | * | 399 |
| northeasterness | 100 | -0.893 | -0.451 | 0.060 | 0.3425 |  | 399 |
| windwardness | 100 | -0.958 | -0.288 | 0.199 | 0.0150 | * | 399 |
| soil_depth | 100 | -0.973 | 0.231 | 0.005 | 0.8475 |  | 399 |
| rockiness | 100 | -0.404 | 0.915 | 0.097 | 0.0425 | * | 399 |
| pH | 100 | -0.886 | 0.463 | 0.164 | 0.0025 | ** | 399 |
| sand | 25 | 0.300 | 0.954 | 0.184 | 0.1300 |  | 99 |
| silt | 25 | -0.525 | -0.851 | 0.208 | 0.1300 |  | 99 |
| clay | 25 | 0.203 | -0.979 | 0.084 | 0.3500 |  | 99 |
| C | 25 | 0.996 | 0.090 | 0.023 | 0.7800 |  | 99 |
| tN | 25 | 0.820 | 0.573 | 0.016 | 0.8600 |  | 99 |
| CN_ratio | 25 | 0.891 | -0.454 | 0.075 | 0.3900 |  | 99 |
| eN | 25 | 0.634 | -0.774 | 0.026 | 0.8100 |  | 99 |
| P | 25 | 0.842 | -0.539 | 0.166 | 0.1000 | . | 99 |
| K | 25 | 0.919 | -0.394 | 0.023 | 0.8100 |  | 99 |
| Ca | 25 | 0.992 | -0.125 | 0.038 | 0.7600 |  | 99 |
| Mg | 25 | 0.662 | -0.750 | 0.148 | 0.2100 |  | 99 |
| Fe | 25 | 0.913 | -0.407 | 0.056 | 0.5600 |  | 99 |
| Mn | 25 | -0.919 | 0.395 | 0.238 | 0.0700 | . | 99 |
| Cu | 25 | -0.761 | 0.648 | 0.273 | 0.0200 | * | 99 |
| Zn | 25 | 0.929 | -0.369 | 0.145 | 0.1700 |  | 99 |
| S | 25 | 0.372 | 0.928 | 0.057 | 0.6300 |  | 99 |
| k | 25 | -0.758 | 0.652 | 0.097 | 0.2900 |  | 99 |

**Table S3.** Checklist of all woody species recorded within Lalashan Forest Dynamics Plot, including Latin species name with authors, Chinese species name (Chinese), family, and species abbreviation (Abbrev.). Abbreviations were created from 4 letters of genus and 4 letters of the the name on the lowest taxonomic level (species, subspecies or variety), with exception of *Chamacyparis formosensis* and *C. obtusa* var. *formosana*, which were abbreviated as Chamform and Chamobtu, respectively.

| Latin species name | Chinese | Family | Abbrev. |
| --- | --- | --- | --- |
| *Acer kawakamii* Koidz. | 尖葉槭 | Aceraceae | Acerkawa |
| *Acer morrisonense* Hayata | 臺灣紅榨槭 | Aceraceae | Acermorr |
| *Acer palmatum* var. *pubescens* Li | 臺灣掌葉槭 | Aceraceae | Acerpube |
| *Barthea barthei* (Hance ex Benth.) Krasser | 深山野牡丹 | Melastomataceae | Bartbart |
| *Cornus kousa* subsp. *chinensis* (Osborn) Q.Y.Xiang | 四照花 | Cornaceae | Bentchin |
| *Berberis hayatana* Mizush. | 早田氏小檗 | Berberidaceae | Berbhaya |
| *Callicarpa randaiensis* Hayata | 巒大紫珠 | Verbenaceae | Callrand |
| *Camellia brevistyla* (Hayata) Cohen-Stuart | 短柱山茶 | Theaceae | Camebrev |
| *Carpinus rankanensis* Hayata | 蘭邯千金榆 | Betulaceae | Carprank |
| *Chamaecyparis formosensis* Matsum. | 紅檜 | Cupressaceae | Chamform |
| *Chamaecyparis obtusa* var. *formosana* (Hayata) Hayata | 臺灣扁柏 | Cupressaceae | Chamobtu |
| *Cleyera japonica* Thunb. | 紅淡比 | Theaceae | Cleyjapo |
| *Daphniphyllum himalaense* (Benth.) Müll.Arg. subsp. *macropodum* (Miq.) T.C.Huang | 薄葉虎皮楠 | Daphniphyllaceae | Daphmacr |
| *Dendropanax dentiger* (Harms) Merr. | 臺灣樹參 | Araliaceae | Denddent |
| *Euonymus spraguei* Hayata | 刺果衛矛 | Celastraceae | Euonspra |
| *Eurya crenatifolia* (Yamam.) Kobuski | 假柃木 | Theaceae | Eurycren |
| *Eurya glaberrima* Hayata | 厚葉柃木 | Theaceae | Euryglab |
| *Eurya loquaiana* Dunn | 細枝柃木 | Theaceae | Euryloqu |
| *Hydrangea angustipetala* Hayata | 狹瓣八仙花 | Saxifragaceae | Hydrangu |
| *Ilex hayatana* Loes. | 早田氏冬青 | Aquifoliaceae | Ilexhaya |
| *Ilex lonicerifolia* Hayata | 忍冬葉冬青 | Aquifoliaceae | Ilexloni |
| *Ilex sugerokii* var. *brevipedunculata* (Maxim.) S.Y. Hu | 太平山冬青 | Aquifoliaceae | Ilexbrev |
| *Ilex suzukii* S.Y. Hu | 鈴木冬青 | Aquifoliaceae | Ilexsuzu |
| *Ilex tugitakayamensis* Sasaki | 雪山冬青 | Aquifoliaceae | Ilextugi |
| *Itea parviflora* Hemsl. | 小花鼠刺 | Saxifragaceae | Iteaparv |
| *Ligustrum liukiuense* Koidz. | 日本女貞 | Oleaceae | Liguliuk |
| *Lindera erythrocarpa* Makino | 鐵釘樹 | Lauraceae | Linderyt |
| *Litsea acuminata* (Blume) Kurata | 長葉木薑子 | Lauraceae | Litsacum |
| *Litsea elongata* var. *mushaensis* (Hayata) J.C. Liao | 霧社木薑子 | Lauraceae | Litsmush |
| *Machilus thunbergii* Siebold & Zucc. | 紅楠 | Lauraceae | Machthun |
| *Michelia compressa* (Maxim.) Sarg. | 烏心石 | Magnoliaceae | Michcomp |
| *Microtropis fokienensis* Dunn | 福建賽衛矛 | Celastraceae | Micrfoki |
| *Neolitsea aciculata* (Blume) Koidz. | 銳葉新木薑子 | Lauraceae | Neolacic |
| *Neolitsea acuminatissima* (Hayata) Kaneh. & Sasaki | 高山新木薑子 | Lauraceae | Neolacum |
| *Osmanthus heterophyllus* (G. Don) P.S. Green | 異葉木犀 | Oleaceae | Osmahete |
| *Photinia niitakayamensis* Hayata | 玉山假沙梨 | Rosaceae | Photniit |
| *Pieris taiwanensis* Hayata | 臺灣馬醉木 | Ericaceae | Piertaiw |
| *Pourthiaea beauverdiana* var. *notabilis* (C.K. Schneid.) Hatus. | 臺灣老葉兒樹 | Rosaceae | Pournota |
| *Pourthiaea villosa* var. *parvifolia* (E. Pritz.) H. Iketani & H. Ohashi | 小葉石楠 | Rosaceae | Pourparv |
| *Prunus phaeosticta* (Hance) Maxim. | 墨點櫻桃 | Rosaceae | Prunphae |
| *Prunus transarisanensis* Hayata | 阿里山櫻花 | Rosaceae | Pruntran |
| *Quercus longinux* Hayata | 錐果櫟 | Fagaceae | Querlong |
| *Quercus sessilifolia* Blume | 毽子櫟 | Fagaceae | Quersess |
| *Quercus stenophylloides* Hayata | 狹葉櫟 | Fagaceae | Quersten |
| *Rhamnus crenata* Siebold & Zucc. | 鈍齒鼠李 | Rhamnaceae | Rhamcren |
| *Rhododendron formosanum* Hemsl. | 臺灣杜鵑 | Ericaceae | Rhodform |
| *Rhododendron leptosanthum* Hayata | 西施花 | Ericaceae | Rhodlept |
| *Rhododendron pseudochrysanthum* Hayata | 玉山杜鵑 | Ericaceae | Rhodpseu |
| *Schima superba* Gard. & Champ. | 木荷 | Theaceae | Schisupe |
| *Skimmia japonica* Thunb. subsp. distincte-venulosa (Hayata) T.C.Ho | 臺灣茵芋 | Rutaceae | Skimvenu |
| *Sorbus randaiensis* (Hayata) Koidz. | 巒大花楸 | Rosaceae | Sorbrand |
| *Styrax formosanus* Matsum. | 烏皮九芎 | Styracaceae | Styrform |
| *Sycopsis sinensis* Oliv. | 水絲梨 | Hamamelidaceae | Sycosine |
| *Symplocos formosana* Brand | 臺灣灰木 | Symplocaceae | Sympform |
| *Symplocos macrostroma* Hayata | 大花灰木 | Symplocaceae | Sympmacr |
| *Symplocos migoi* Nagarn. | 擬日本灰木 | Symplocaceae | Sympmigo |
| *Tetradium glabrifolium* (Champ. ex Benth.) T.G. Hartley | 賊仔樹 | Rutaceae | Tetrglab |
| *Tetradium ruticarpum* (A. Juss.) T.G. Hartley | 吳茱萸 | Rutaceae | Tetrruti |
| *Trochodendron aralioides* Siebold & Zucc. | 昆欄樹 | Trochodendraceae | Trocaral |
| *Tsuga chinensis* var. *formosana* (Hayata) H.L. Li & H. Keng | 臺灣鐵杉 | Pinaceae | Tsugform |
| *Vaccinium bracteatum* Thunb. | 米飯花 | Ericaceae | Vaccbrac |
| *Viburnum foetidum* var. *rectangulatum* Rehder | 狹葉莢蒾 | Caprifoliaceae | Viburect |
| *Viburnum luzonicum* Rolfe | 呂宋莢蒾 | Caprifoliaceae | Vibuluzo |
| *Viburnum sympodiale* Graebn. | 假繡球 | Caprifoliaceae | Vibusymp |
| *Viburnum urceolatum* Siebold & Zucc. | 壺花莢蒾 | Caprifoliaceae | Vibuurce |
